# Changes in the plasmodesma structure and permeability at the bundle sheath and mesophyll interface during the maize C4 leaf development

**DOI:** 10.1101/2020.09.30.320283

**Authors:** Peng Gao, Baijuan Du, Pinghua Li, Byung-Ho Kang

**Author notes:** Author for correspondence: Byung-Ho Kang (, 852-3943-6101). Author contributions: GP and BHK: Planned and designed the research. GP: Performed microscopy imaging, immunolocalization, and qRT-PCR. BD and PL: Maintained maize mutant lines, carried out transcriptomic data analysis and gene expression correlation test. GP, BD, PL, and BHK: Interpreted the data. GP and BHK: Wrote the manuscript.

## Abstract

Plasmodesmata are intercellular channels that facilitate molecular diffusion between neighboring plant cells. The development and functions of plasmodesmata are controlled by multiple intra- and intercellular signaling pathways. Plasmodesmata are critical for dual-cell C4 photosynthesis in maize because plasmodesmata at the mesophyll and bundle sheath interface mediate exchange of CO_2_-carrying organic acids. We examined developmental profiles of plasmodesmata and chloroplasts in the maize leaf from young cells in the base to mature cell in the tip using microscopy approaches. Young mesophyll and bundle sheath cells in the leaf base had proplastids, and their plasmodesmata were simple, devoid of cytoplasmic sleeves. In maturing cells where Kranz anatomy and dimorphic chloroplasts were evident, we observed extensive remodeling of plasmodesmata that included acquisition of an electron-dense ring on the mesophyll side and cytoplasmic sleeves on the bundle sheath side. Interestingly, the changes in plasmodesmata involved a drop in symplastic dye mobility and suberin accumulation in the cell wall, implying a more stringent mesophyll-bundle sheath transport. We compared kinetics of the plasmodesmata and the cell wall modification in wildtype leaves with leaves from *ppdk and dct2* mutants with defective C4 pathways. Plasmodesmata development, symplastic transport inhibition, and cell wall suberization were accelerated in the mutant lines, probably due to the aberrant C4 cycle. Transcriptomic analyses of the mutants confirmed the expedited changes in the cell wall. Our results suggest that a regulatory machinery at the mesophyll-bundle sheath boundary suppresses erroneous flux of C4 metabolites in the maize leaf.

**Significance Statement:** Plasmodesmata in the maize Kranz anatomy mediate the exchange of organic acids between mesophyll and bundle sheath. Since solute diffusion through plasmodesmata is governed by solute concentration gradients, a balanced distribution of C4 metabolites is critical for concentration of CO_2_ in the bundle sheath. Plasmodesmata bridging the mesophyll and bundle sheath cytoplasm have a cylindrical cavity, which can facilitate molecular movements, and a valve-like attachment. Construction of the sophisticated plasmodesmata was linked to C4 photosynthesis, and plasmodesmata assembly finished more rapidly in maize mutants with defective C4 pathways than in wild-type plants. These results suggest that the specialized plasmodesmata contribute to controlled transport of C4 metabolites.

## Introduction

C4 photosynthesis involves a biochemical cycle that pumps CO_2_ to curb energetically wasteful photorespiration (1, 2). The cycles require two compartments, one for producing four carbon (C4) acids through fixation of atmospheric CO_2_ and the other for decarboxylating the C4 acids to provide Rubisco and CO_2_. The Kranz anatomy is the leaf architecture that partitions the two compartments into two cell types (3, 4).

The classical NADP-malic enzyme-mediated C4 pathway occurs in the maize leaf. In the leaf, bundle sheath (BS) cells form a ring around vascular cells and mesophyll (M) cells are located outside the BS cells (5). Phosphoenolpyruvate carboxylase in the M cell cytosol captures CO_2_ to produce oxaloacetate, which is subsequently reduced to malate in the M chloroplast. Malate moves to the BS cell and is subsequently imported into the BS chloroplast by DCT2, a dicarboxylic acid transporter. Decarboxylation of malate produces pyruvate and CO_2_ and pyruvate return to the M cell. The pyruvate orthophosphate dikinase PPDK regenerates phosphoenolpyruvate from pyruvate in the M chloroplast for another round of CO_2_ pumping (6). Maize mutant lines in which genes encoding DCT2 or PPDK are inactivated have been characterized. (7, 8). C4 photosynthesis is defective in leaves of the mutant plants, but their Kranz anatomy is not compromised.

The M and BS chloroplasts of the maize have different energy requirements and have different ultrastructural features (9). In the M chloroplast, both photosystem II (PSII) and photosystem I (PSI) mediate linear electron flow to produce ATP and NADPH. The BS chloroplast produces ATP via PSI-mediated cyclic electron flow since reducing power comes with malate that originates in M cells. To accommodate PSII and PSI complexes, the M chloroplast have grana stacks and stroma lamellae. Thylakoids in BS chloroplasts consist of unstacked thylakoids because they lack PSII (10–12). Proplastids in immature cells at the maize leaf base are morphologically similar, indicating that the dimorphic chloroplasts arise as the two cell types mature along the leaf developmental gradient toward the tip (13).

Plasmodesmata (PD) are channels through the cell wall that connect the cytoplasms of neighboring plant cells. Transmission electron microscopy (TEM) studies have revealed that each channel consists of the plasma membrane and a membrane tubule called the desmotubule that originates from the endoplasmic reticulum (14, 15). Metabolites, signaling molecules, and small proteins can diffuse into adjacent cells through the cytoplasmic sleeve, the gap between the plasma membrane and the desmotubule (16–18). Callose, a β-1,3-glucan, is associated with PD, and increases in callose amounts cause constriction of the PD to inhibit diffusion (19, 20). PD are assembled in the cell plate during cytokinesis, and the nascent PD are morphologically simple. As the cells that they connect differentiate, the PD evolve, and their structures alter (14). PD connecting mature companion cells and sieve elements have a tree-like shape, highly branched on the companion side. PD density in a plant tissue also varies depending on the rate of symplastic flux in the tissue.

The PD at the M-BS interface in the maize leaf mediate the transport of molecular components of the C4 cycle (21, 22). The movement of malate to the BS cell and recycling of pyruvate to the M cell are dependent on passive transport (23). Transamination of oxaloacetate generates aspartate in the M cytosol, and this C4 acid also enters BS through PD (24, 25). The BS cell wall is suberized to block leakage of CO_2_ and sucrose and to transport organic acids exclusively through PD (26). PD that transverse the M-BS cell walls occur in large clusters in the maize leaf. The PD density in the maize leaf is approximately 9 times higher than in the rice leaf, a C3 crop plant. The high degree of symplastic connectivity of the PD in maize reflects its significance in solute exchange in the C4 system (27). It has been shown that PD linking M and BS cells have a sphincter on the M side in dual-cell C4 plants including maize and sugarcane (28, 29).

Despite the pivotal role of PD in the maize C4 photosynthesis, how the symplastic transport machinery in the Kranz anatomy is assembled and regulated had not been established. Using a suite of microscopy methods combined with transcriptomic analyses, we characterized the ontogeny and permeability of PD at the M-BS boundary in association with dimorphic chloroplast development in the maize leaf. We provide evidence for a link between PD gating and proper operation of the C4 pathway.

## Results

### Development of dimorphic chloroplasts in the maize leaf

We examined chloroplast structures in the maize leaf with TEM and electron tomography (ET) to characterize the ultrastructural dynamics involved in the C4 differentiation. The four locations along the developmental gradient of third leaves of 9-day-old maize seedlings were selected as defined in Li *et al*. (9) (Fig. 1A). The location of ligule 2 in leaf 3 was set as the reference point (0 cm), and maize leaves were cut perpendicular to the leaf axis at −4 cm, 0 cm, 4 cm, and 10 cm (Fig. 1B). Thin cross sections (0.2-0.3 mm) were isolated and preserved by high-pressure freezing and freeze-substitution for ET analyses. For consistency in comparing the four stages among genotypes, our examination was restricted to minor veins because they outnumber midveins and intermediate veins by 7 to 10 fold (28). We examined intermediate veins in the −4-cm samples, because minor veins do not extend to the young leaf tissue, and the cell organization of intermediate veins is more similar to that of minor veins than that of midveins. Intermediate and minor veins correspond to rank 1 and rank 2 intermediate veins, respectively, according to the classification by Sedelnikova *et al*. (5).

**Fig. 1.**
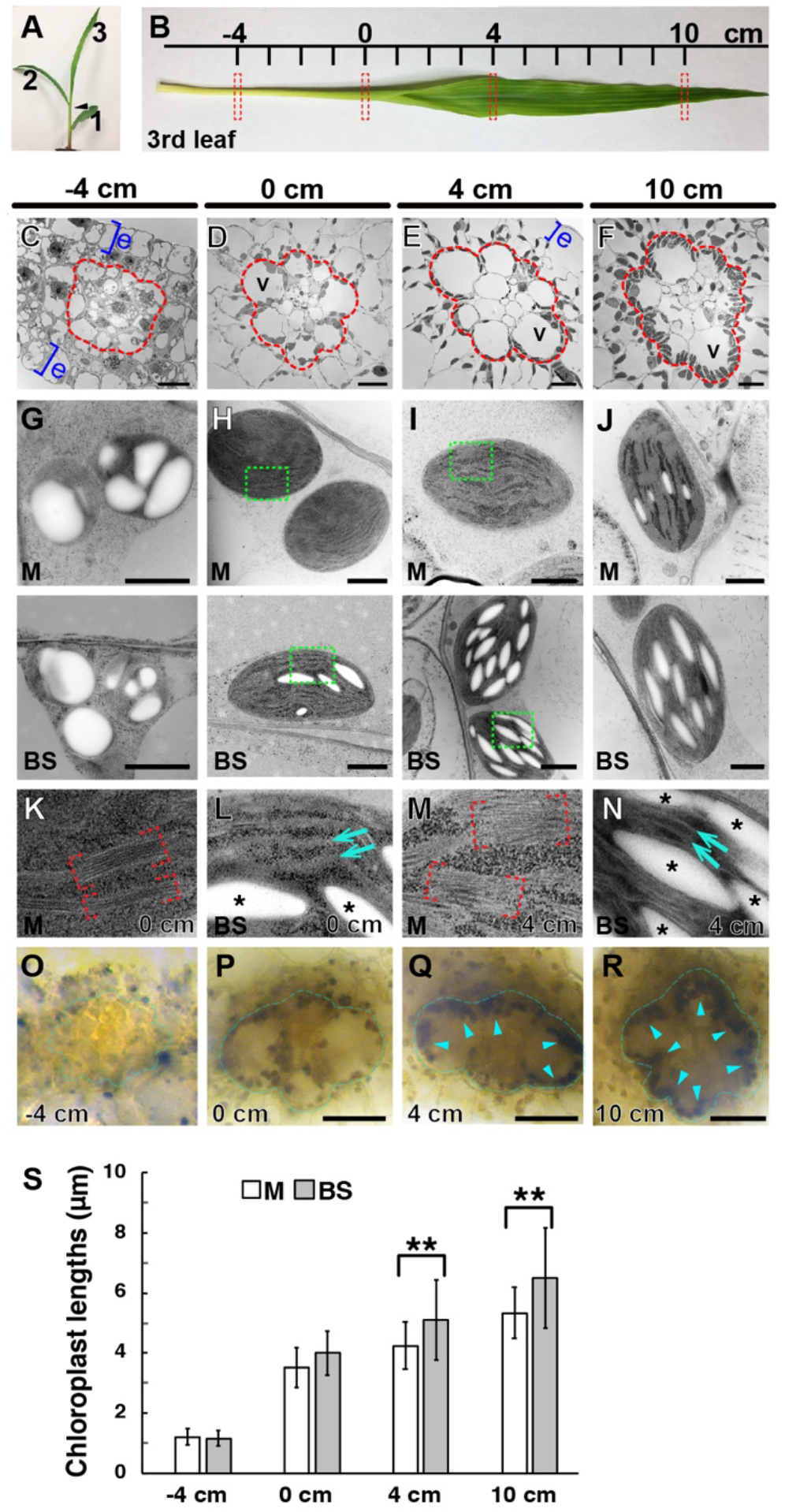
Analysis of development of dimorphic chloroplasts in the B73 seedling leaf using TEM. **(A)** A 9-day old maize B73 seedling. The first (1), second (2), and third (3) leaves are marked. **(B)** The four positions (−4, 0, 4, 10 cm) of the third leaf were isolated for microscopy analysis (dashed rectangles). **(C-F)** Low-mag TEM images of the four segments in B. The vascular bundle and BS cells are enclosed in the red outline in each panel. Blue brackets indicate epidermis (e) layers. V: vacuole. Scale bars: 10 μm. **(G-J)** TEM photos of chloroplasts in M cells (upper row) and BS cells (lower row) in the four locations along the leaf. Scale bars: 1.0 μm. **(K-N)** Higher magnification micrographs of the boxed areas in H and I to show grana stacks in M chloroplasts (red brackets in K and M) and stroma lamellae in BS chloroplasts (arrows in L and N). Starch particles are denoted with asterisks. **(O-R)** Starch granules after iodine staining. Starch accumulation in BS cells (outlined with blue lines) is discerned in the 4- and 10-cm samples (blue arrowheads). Scale bars: 50 μm. **(S)** Lengths of M (white bars) and BS (grey bars) chloroplasts in TEM micrographs. Longer axes of chloroplasts (n = ~25 for each stage) were measured. Error bars correspond to standard deviations (SD) (**\*\***, p<0.01; Student’s t-test).

Samples were examined by low- (Fig. 1C-F) and high-magnification (Fig. 1G-J) TEM. We were able to discern the M and BS cells in the −4-cm cross section but the concentric arrangement indicative of Kranz anatomy was not apparent (Fig. 1C). In 0-cm cross sections, however, the vascular bundle, BS cells, and M cells outside the BS layer were visible (Fig. 1D). Proplastids in M and BS cells at −4 cm were indistinguishable: They were round with average diameters of 1.5-1.6 μm, and starch particles occupied most of their stroma (Fig. 1G and O) as reported in Majeran *et al*. (13). Chloroplasts of the two cell types in the 0-cm zone were larger and had more thylakoids than those of the −4-cm zone. M chloroplasts had grana stacks but few starch particles, whereas BS chloroplasts had starch particles but grana stacks were rare, displaying signs of differentiation (Fig. 1H, K, and L). The differing thylakoid morphologies of the M and BS cells in the 0-cm sections indicate that the two cell types express different sets of chloroplast proteins.

The chloroplast dimorphism of Kranz anatomy was clearly observed in 4-cm zone chloroplasts, (Fig. 1G-J). Grana stacks were abundant in M chloroplasts, but thylakoids were mostly unstacked in BS chloroplasts (Fig. 1I, M, and N). Thylakoid architectures in the two cell types in the 4-cm sections persisted in 10-cm sections (Fig. 1J). When starch particles were stained with iodine, darkly stained particles accumulated in BS cells of leaf sections from 4-cm and 10-cm positions (Fig. 1O-R). The starch accumulation correlated with the extent of chloroplast development (Fig. 1K-N). Chloroplasts were ovoid in both cell types, but BS chloroplasts were more stretched out than M chloroplasts (Fig. 1S).

To delineate thylakoid assembly more accurately, we carried out ET analysis of chloroplasts in −4-, 0-, and 4-cm leaf sections. At −4 cm, plastids had simple thylakoid networks, and grana stacks consisted of two or three layers in both cell types (Fig. 2A and B). In M chloroplasts in 0-cm sections, each granum had approximately five layers on the average, and grana stacks densely populated the stroma (Fig. 2C). Thylakoids in BS chloroplasts of the same stage had extended stoma lamellae and grana stacks consisting of 2-4 disks (Fig. 2D). The thylakoid structures in M chloroplasts were unambiguously distinct from those in BS chloroplasts at 4 cm (Fig. 2E and F). Grana stacks in M chloroplasts often had more than 10 disks interconnected by helical tubules. By contrast, thylakoids BS chloroplasts were composed of unstacked lamellae. Grana stacks were rare, and those observed had only two or three layers (Fig. 2G), indicating that stroma lamellae expanded and grana stacks shrank during leaf maturation. Moreover, grana stacks in BS chloroplasts at 4 cm were smaller than those in younger cells (Fig. 2H).

**Fig. 2.**
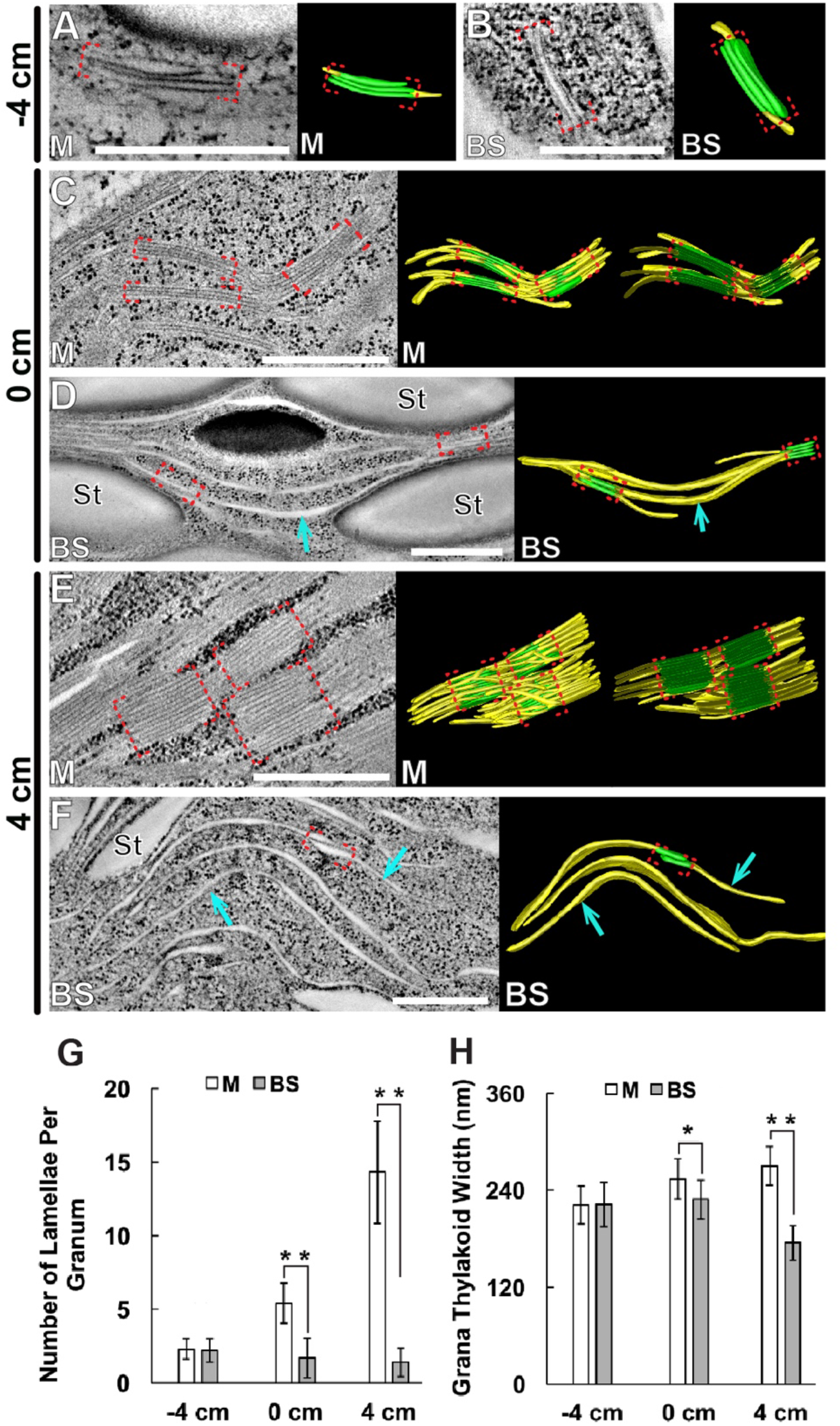
ET analysis of thylakoid assembly in chloroplasts of the maize leaf. **(A-F)** Tomographic slice images of thylakoids in M and BS chloroplasts (left) and 3D models (right) in (A and B) −4-cm, (C and D) 0-cm, and (E and F) 4-cm sections. Stacked and unstacked regions of thylakoids are colored in green and yellow, respectively in 3D models. Grana stacks were highlighted with red brackets. In panels C and E, the thylakoid models were clipped to reveal their stack architectures. Scale bars: 500 nm. **(G)** Average numbers of disks per granum in M and BS chloroplasts (n=10 per cell type and stage). Error bars depict standard errors (SEM) (**\*\***, p<0.01; ns, not significant; Student’s t-test). **(H)** Average widths of grana stacks in M and BS chloroplasts (n=10 per cell type and stage). Error bars depict SEM (**\***, p<0.05; **\*\***, p<0.01; Student’s t-test).

### Localization of chloroplast proteins in M and BS cells by immunofluorescence

Our ET results indicated that the chloroplast structures in M and BS deviated from each other in the 0-cm location and were fully differentiated by 4 cm (Figs. 1 and 2). We performed immunofluorescence localization of subunits in the thylakoid membrane protein complexes and of Rubisco to compare the structural dimorphism with macromolecular compositions of the chloroplasts. We utilized antibodies against PsbO (a subunit of PSII), Lhca (a subunit of the light harvesting complex of PSI), CURT1A, (a protein that stabilizes the grana margin), and the large subunit of Rubisco. None of these proteins were detected in the −4-cm samples (Fig. 3A, D, G, and J). In 0-cm samples, we were able to image fluorescence from both M and BS chloroplasts, but significant differences between the two cell types were not detected (Fig. 3B, E, H, and K).

**Fig. 3.**
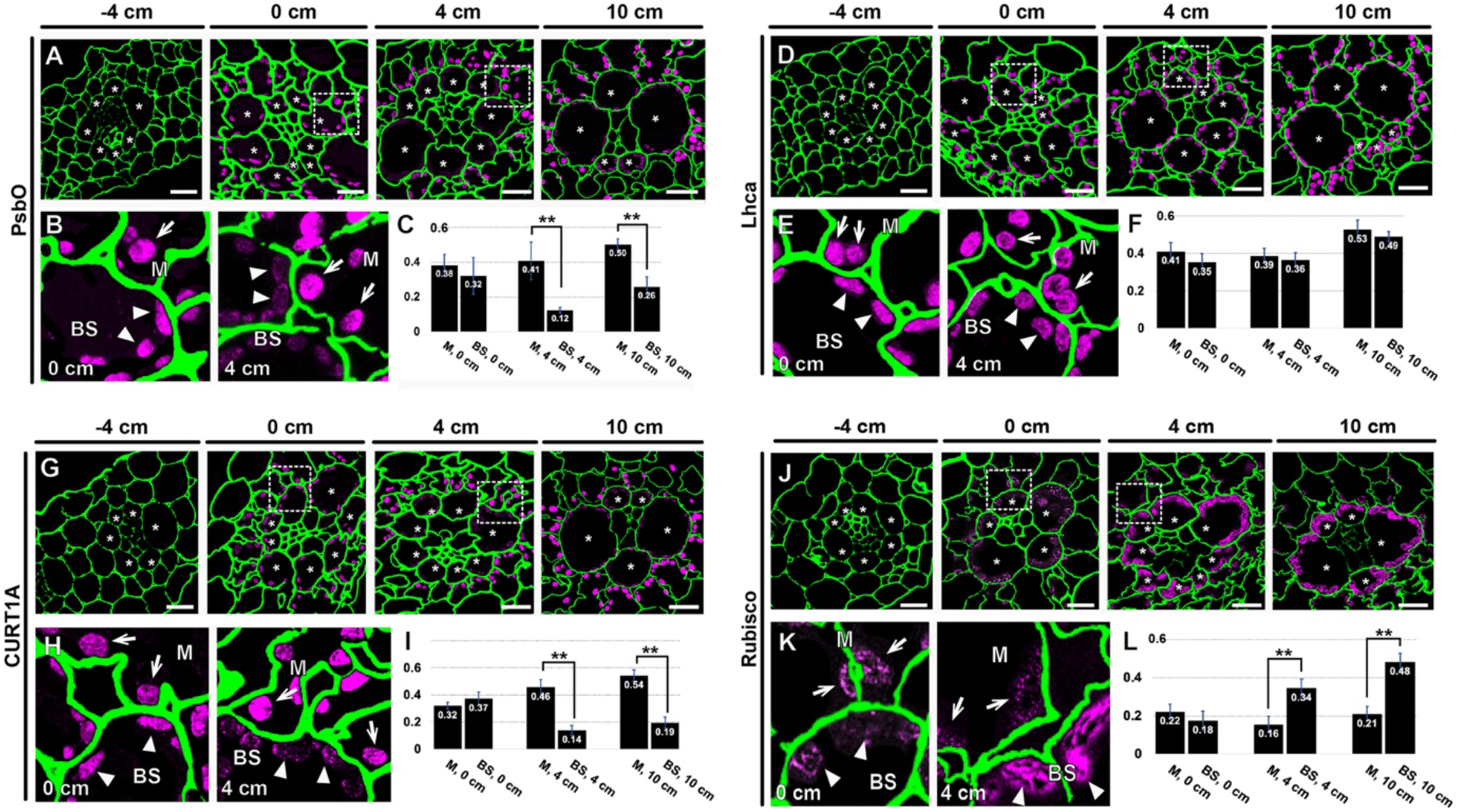
Immunofluorescence localization of chloroplast proteins in M and BS cells at the four leaf development stages. Localization of (**A-C**) PsbO, (**D-F**) Lhca, (**G-I**) CURT1A, and (**J-L**) Rubisco small subunit was detected. For each protein, low magnification micrographs (upper mages) and higher magnification micrographs of the boxed areas in 0 and 4 cm sections (lower left images) are shown. Cell walls (green) were stained to illustrate Kranz anatomy. BS cells are marked with asterisks in the low magnification images. M and BS chloroplasts are indicated with arrows and arrowheads, respectively, in high magnification images. M chloroplasts are more strongly stained by the PsbO and CURT1A antibodies in 4-cm section (panels B and H, respectively). The opposite is true Rubisco in 4-cm sections (panel K). Scale bars: 10 μm. Inset graphs (panels C, F, I, and L) plot fluorescence intensities from M and BS chloroplasts in 0- 4-, and 10-cm sections. Error bars depict standard SD (**\*\***, p<0.01; Student t-test).

Differential enrichment of PsbO, CURT1A, and the Rubisco large subunit in either M and BS chloroplasts was clearly seen in 4-cm samples (Fig. 3B, H, and K). PSII and its light harvesting complexes contribute to the formation of grana stacks and CURT1A stabilizes the grana margin (10, 30). Consistent with the proliferation and removal of grana stacks in M and BS chloroplasts, respectively, PsbO and CURT1A were detected exclusively in M chloroplasts in the 4-cm regions (Fig. 3B, C, H, and I). Lhca-specific fluorescence, indicative of PSI, was detected in both M and BS chloroplasts throughout the leaf developmental gradient (Fig. 3E and F). Significantly more signal due to the Rubisco subunit was detected in BS chloroplasts than in M chloroplasts in the 4-cm section (Fig. 3K and L). The specific enrichment of Rubsico suggests that the Calvin cycle operates primarily in BS chloroplasts in the 4-cm leaf cells.

Chlorophyll fluorescence from PSI is enriched in the spectral window above 700 nm, whereas that from PSII dominates the window below 700 nm; this feature has been utilized to visualize dimorphic chloroplasts in C4 plants (31, 32). We therefore tested whether PSII fluorescence is weaker from BS in the 4-cm leaf samples. PSII intensities were similar in M and BS cells in the −4- and 0-cm sections (SI Appendix, Fig. S1A and B). By contrast, PSII-specific fluorescence was lower in BS cells than M cells in 4-and 10-cm sections (SI Appendix, Fig. S1C and D). There were no difference in the fluorescence detection range of 720-800 nm (*i.e*., PSI) in chloroplasts from BS cells compared to those from M cells at any stage. Thus, the PSI- and PSII-specific chlorophyll fluorescence imaging agrees with the immunofluorescence localization data.

### Development of PD in the maize leaf

It was previously shown that PD connecting an M and BS cell pair has a collar around its desmotubule, termed a sphincter; the sphincter is located on the M side (22). In TEM micrographs of −4-cm leaf sections, PD were simple and devoid of sphincters. Thickness of the cell wall between M and BS cells almost doubled from the −4-cm section to the 0-cm section (Fig. 4 and SI Appendix, Fig. S2). As the cell wall thickened, PD stretched out, but no significant ultrastructural changes were seen. Electron-dense collars were observed in PD of 4-cm and 10-cm leaf samples (Fig. 4A). When visualized with ET, the doughnut-shaped sphincters appear to surround the desmotubule and press the plasma membrane outward, enlarging the neck region. Cytoplasmic sleeves were observed in the BS half of PD in 4-cm sections as were sphincters in the M side (Fig. 4B and C). These structural changes in PD correlated with chloroplast differentiation between 0-cm and 4-cm.

**Fig. 4.**
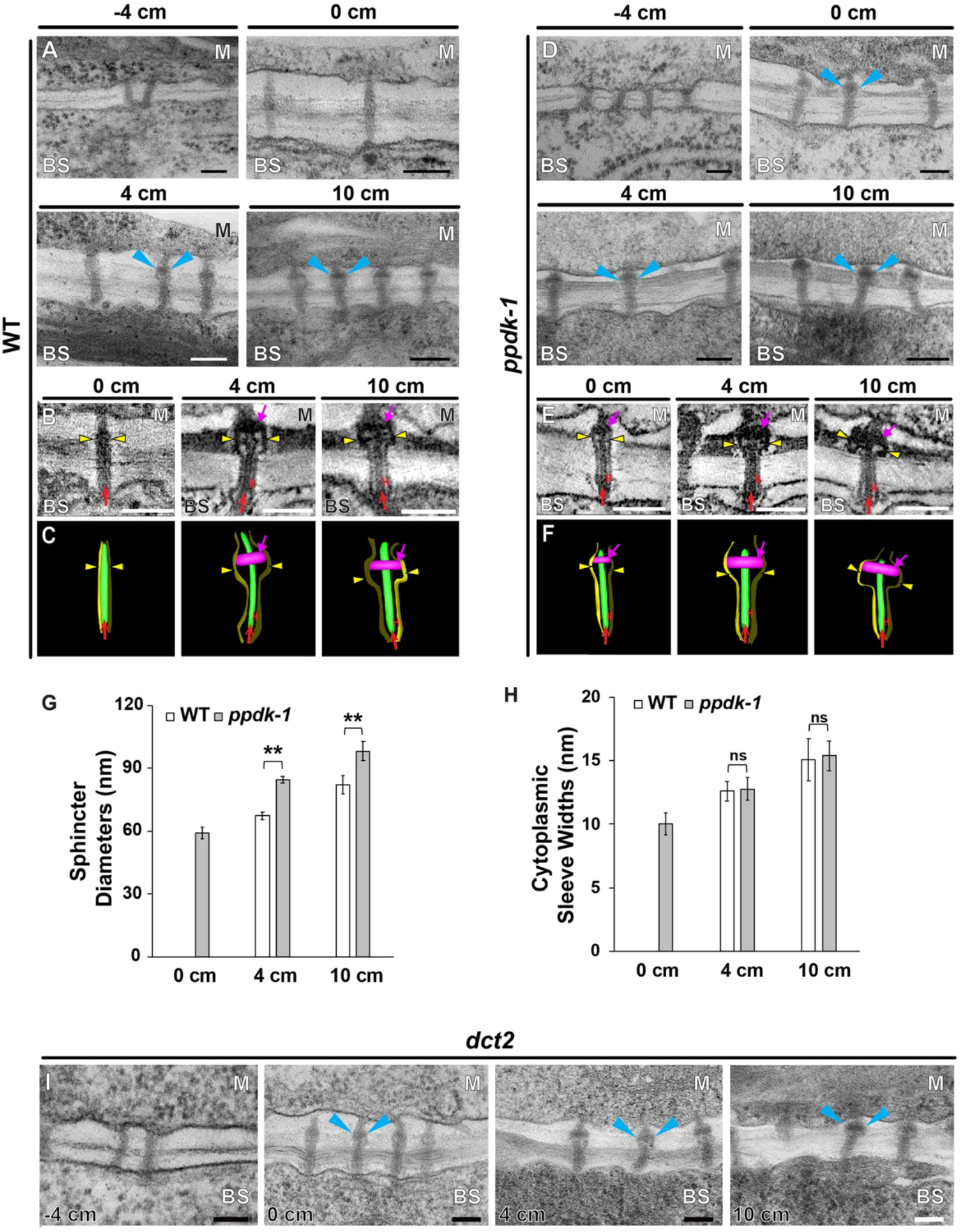
Analyses of PD that connect M and BS cells. **(A)** TEM micrographs of PD at the M-BS interface in wild-type B73 (WT) leaves. PD sphincters are marked with arrowheads. Scale bars, 100 nm. **(B and C)** PD and the plasma membrane at the M-BS interface as observed in (B) ET slice images and (C) 3D models. Sphincters, desmotubules, plasma membrane, and cytoplasmic sleeves are marked with magenta arrows, red arrows, yellow arrowheads, and red “H”s respectively. Scale bars, 150 nm. **(D and F)** TEM micrographs of PD at the M-BS interface in (D) *ppdk-1* leaves, (E) ET slice images, and (F) 3D models. Sphincters, desmotubules, plasma membrane, and cytoplasmic sleeves are marked with magenta arrows, red arrows, yellow arrowheads, and red “H”s respectively. (**G and H**) Morphometric comparisons of PD at the M-BS interface in WT (white bars) and *ppdk-1* (gray bars) leaves. (G) Sphincter diameters and (H) cytoplasmic sleeve widths were measured from tomograms at the three stages. Approximately 20 PD from three different plants were examined for each genotype and each stage. Error bars depict SEM (**\*\***, p<0.01; ns: not significant; Student’s t-test). **(I)** TEM micrographs of PD at the M-BS interface in *dct2* leaves. PD sphincters were discerned in 0-cm sections from the three mutant leaves. Sphincters are indicated by arrowheads.

We then examined PD that join the same cell types to determine whether they undergo changes similar to those observed for PD that join M and BS cells. The sphincters were discerned in PD that connect two M cells in 4- and 10-cm sections. However, sphincters were not observed in PD connecting two BS cells (SI Appendix, Fig. S2). The PD connecting two M cell walls had sphincter collars on both sides, indicating that the attachment is derived from the M cell. The thickness of the BS cell wall increased from −4 to 4 cm, but the M cell wall stopped thickening at 0 cm (SI Appendix, Fig. S2).

Because biogenesis of dimorphic chloroplasts is essential for C4 photosynthesis, the PD remodeling might be related to the C4 cycle. To test this notion, we compared PD in *ppdk-1* and *dct2* maize plants in which key enzymes of C4 cycle are mutated. PD developed more rapidly in the maize mutant lines than in wild-type plants (Fig. 4D-J). PD in the *ppdk-1* and the *dct2* mutant leaves elongated as the cell wall thickened from −4 to 0 cm. ET imaging revealed that sphincter assembly had begun before 0 cm in the mutant leaves (Fig. 4D, I, and J). The PD in the 0-cm mutant leaves had acquired cytoplasmic sleeves. These morphological changes were not detected in wild-type PDs until 4 cm (Fig. 4A-C). Sphincter sizes were larger in *ppdk-1* PD than wild-type PD at the same leaf positions (Fig. 4G). However, widths of cytoplasmic sleeves did not differ between mutant and wild-type plants (Fig. 4H). The accelerated PD changes were observed in the *dct2* mutant (Fig. 4I) and two other *ppdk* mutant alleles (SI Appendix, Fig. S3).

Maize plants homozygous for *ppdk-1 and dct2* mutations are seedling lethal, but their development from germination through 9 days of growth (*i.e*., the time of sampling) was relatively normal (SI Appendix, Fig. S3). The Kranz anatomy was not affected by the deficiency in C4 pathways, although their leaves were pale green. Chloroplast proteins were differentially enriched in M and BS chloroplasts between the 0-cm and 4-cm stages in the mutant leaves as in the wild-type leaf (SI Appendix, Fig. S5). However, BS chloroplasts in the mutants had fewer starch particles than those in wild-type leaves (SI Appendix. Fig. S6). Transcriptomic datasets from *ppdk-1* and *dct2* mutant leaves are available and we performed gene expression correlation test using the RNA-seq datasets (7, 8). In these studies, leaf specimens using the same sampling design as used here. Correlation coefficients of genes were calculated from their expression profiles at the four leaf positions in wild type and the mutant lines. The mutations did not cause large scale transcriptomic alterations in mutant leaves (SI Appendix, Fig. S4).

### PD remodeling is accompanied by reduced symplastic dye movement between MS and BS cells

To investigate consequences of the PD remodeling on transport capacity, we carried out a carboxyfluorescein-diacetate (CFDA) movement assay. After leaves were fed with CFDA solution from their base, cross sections (1-2 mm thick) at 0, 4, and 10 cm were dissected. In 0-cm sections of wild-type leaves, both M and BS cells had CFDA fluorescence (Fig. 5A and B). By contrast, the fluorescence was highly confined to BS cells in 4- and 10-cm samples (Fig. 5A and B). These results indicate that PD permeability is reduced in the older leaf tissues. In *ppdk-1* and *dct2* mutant leaves, CFDA was limited to BS cells in all three locations (Fig. 5C and D). CFDA was detected in M cells of wild-type and mutant leaves in all three stages when the dye solution also contained 2-deoxy-D-glucose (DDG), a callose synthase inhibitor (Fig. 5E-G), indicating that callose synthesis is required for the block in dye transport. When we calculated the ratios of fluorescence intensities in M-BS cell pairs, the largest differences were detected in 0-cm samples from wild-type and mutant leaves because the dye failed to diffuse into M cells from BS cells at this stage (Fig. 5B, C, D, and H). We concluded that C4 development in the maize leaf involves a decrease in symplastic permeability between M and BS cells and that the decrease is temporally associated with structural changes in PD.

**Fig. 5.**
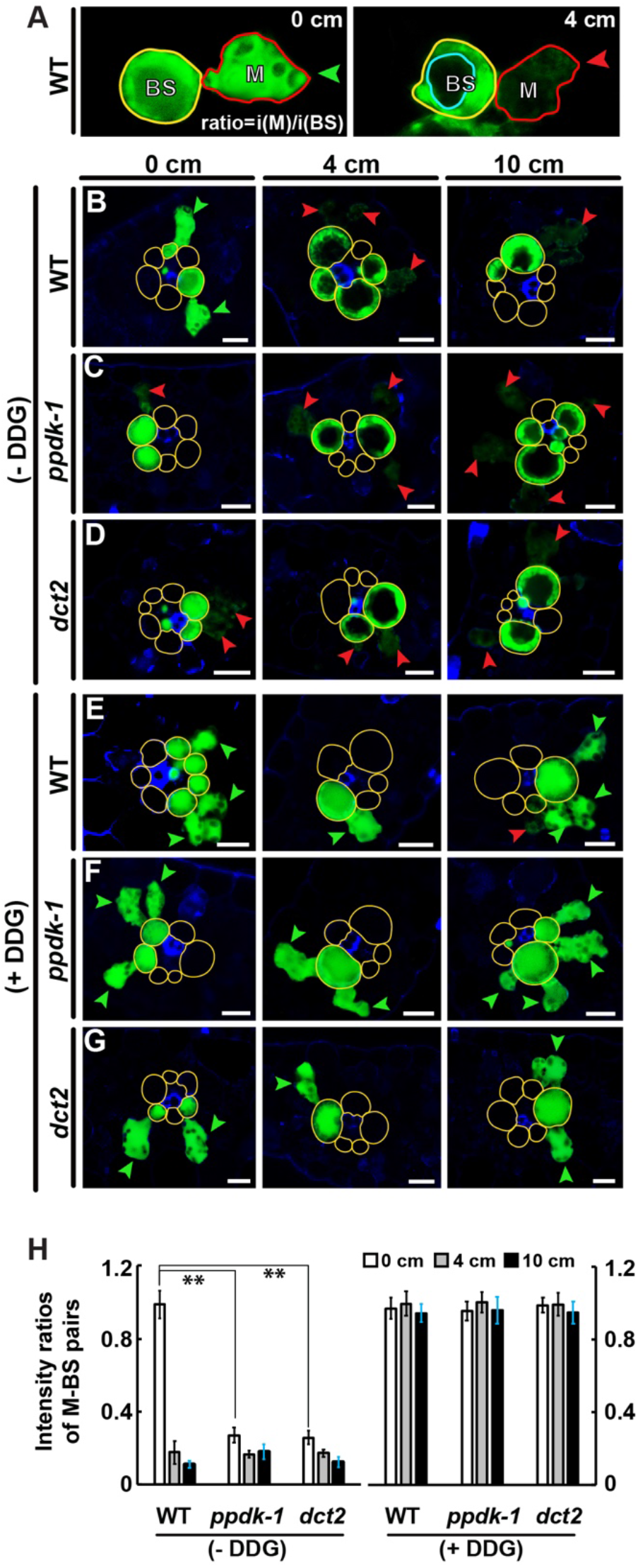
CFDA movement assay. **(A)** Confocal laser scanning micrographs of CFDA fluorescence (green) in WT leaf cross sections (0 and 4 cm). Typical M-BS pairs at 0 cm (left) and 4 cm (right) in WT leaf cross sections emit CFDA fluorescence. Green arrowhead marks fluorescence in M cell comparable to that in BS cell. Red arrowheads indicate faint fluorescence in M cell. Light blue compartment indicates central vacuole. **(B-G)** WT, *ppdk-1*, and *dct2* leaves (B-D) without DDG or (E-G) with DDG. M-BS units in which the M cell display fluorescence comparable to (less than 30% reduction) the BS cell are denoted with green arrowheads. Red arrowheads mark M cells in which fluorescence is weaker (less than 30% that in BS cell). Note that M cells in the 0-cm *ppdk-1* and *dct2* sections do not contain dye. BS cell walls are highlighted with yellow lines. Scale bars: 25 μm. **(H)** CFDA fluorescence ratios in M-BS units. The fluorescence intensities from the cytosols of M and BS cells were quantified to determine ratios. Central vacuoles were excluded when intensities were calculated (**\*\***, p<0.01; Student’s t-test).

### Suberin deposition in the BS cell wall is linked to PD remodeling

The BS wall accumulates suberin, and the hydrophobic stratum blocks apoplastic diffusion of photosynthetic metabolites (26). Since this barrier could restrict solute exchange between M and BS exclusively through PD, we assayed for suberin deposition in the cell wall of the four-leaf zones. The BS cell wall in 0-cm cross sections did not exhibit fluorescence. Only xylem cells in the vascular bundle were stained in the sections (Fig. 6A-C). BS cell walls in 4-cm sections were positive for suberin staining, and the fluorescent outline visualized the BS cell layer outside the vascular bundle (Fig. 6A-C). By contrast, suberin was clearly observed in BS cell walls of 0-cm leaf sections from *ppdk-1* and *dct2* plants (Fig. 6D-I). The staining became stronger at later leaf developmental stages of wild-type as well as the mutants (Fig. 6J).

**Fig. 6.**
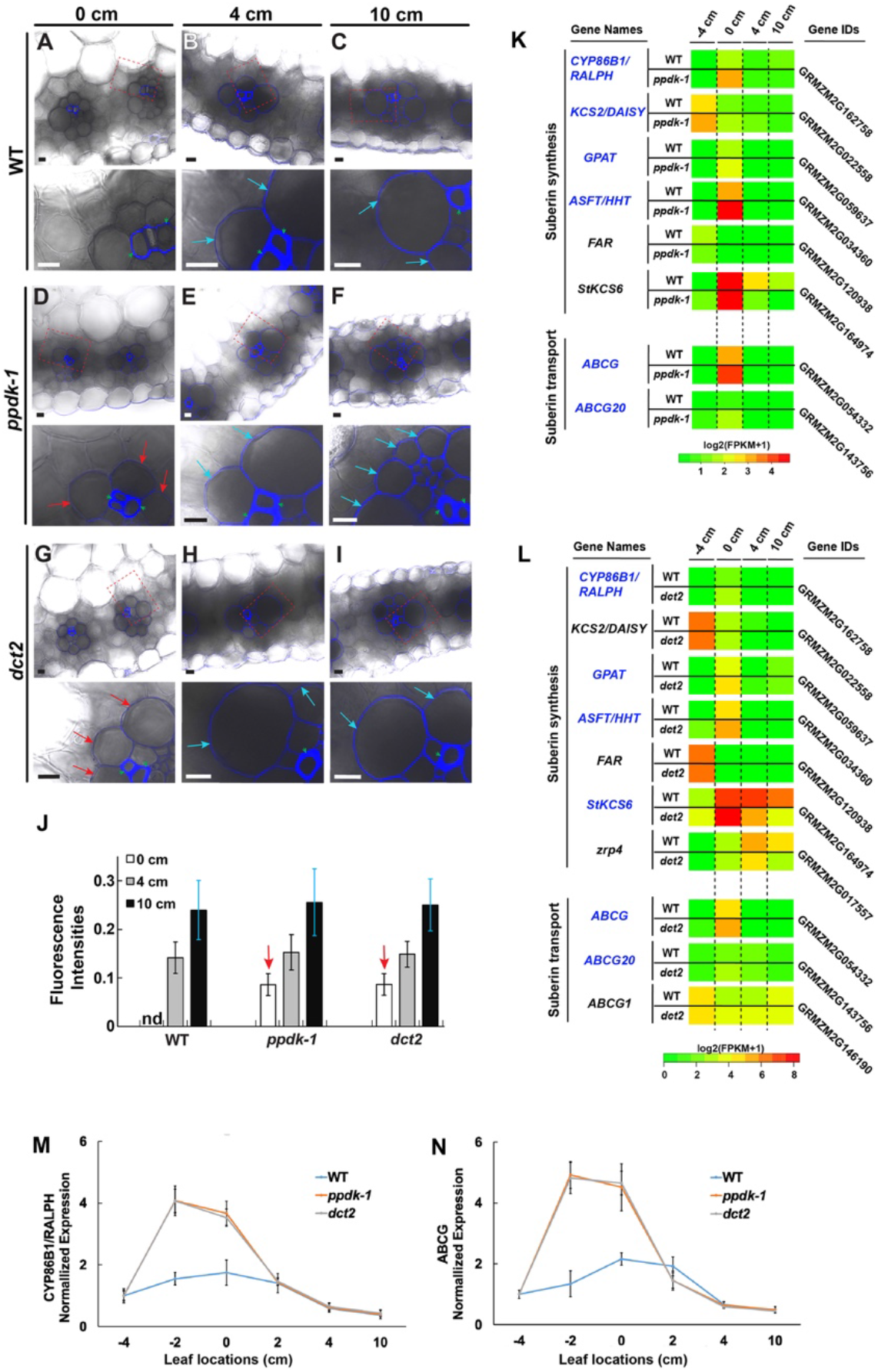
Suberin accumulation in BS cell wall. **(A-I)** Leaf sections at 0 cm, 4 cm, and 10 cm from (A-C) WT B73, (D-F)*ppdk-1*, and (G-I) *dct2* plants stained for suberin. In each panel, lower micrograph images correspond to higher magnifications of the boxed areas in upper micrographs. BS and xylem cell walls visualized by suberin staining are indicated with arrows (red arrows for 0-cm and light blue arrows for 4- and 10-cm images) and arrowheads, respectively. Note that BS cell walls in 0-cm *ppdk-1* and *dct2* samples stain for suberin staining (D and G), whereas BS cell walls in 0-cm WT sample do not (A). Scale bars: 10 μm. **(J)** Fluorescence intensities of BS cell walls in WT, *ppdk-1*, and *dct2* leaf 0-cm (white bars), 4-cm (gray bars), and 10-cm (black bars) sections. Fluorescence values of BS cell walls were normalized to those of adjacent xylem walls. **(K and L)** Heat maps illustrating changes in transcript levels of genes involved in suberin synthesis or transport in *ppdk-1* (K) and *dct2* (L) leaves. All genes with FPKM values higher than 1 were included in the heat maps (SI Appendix, Dataset S1). Genes transcribed more actively in the mutant than WT in 0-cm are in blue letters. **(M and N)** qRT-PCR-based quantification of (M) *CYP86B1* and (N) *ABCG* in six locations in the maize leaf. *GAPDH* was used as the reference gene. Error bars indicate SD.

We examined expression levels of genes that play roles in suberin synthesis or export in the RNA-seq datasets from *ppdk-1* and *dct2* (7, 8). Maize genes associated with cell wall suberization were identified with the MapMan tool for functional classification (http://gabi.rzpd.de/projects/MapMan/). Most of the identified genes exhibited strong transcriptional activities in the −4- or 0-cm zones (Fig. 6K and L). When transcript levels in wild-type and the mutant lines were compared, suberin-related genes were more upregulated in the *ppdk-1* (6 out of 8) and *dct2* (7 out of 10) leaves than wild-type leaves, in agreement with the suberin staining results. To evaluate temporal expression profiles more accurately, we carried out qRT-PCR analysis of *CYP68B* (also known as *RALPH*; GRMZM2G162758) and *ABCG* (GRMZM2G054332) genes in six leaf regions. Transcripts from the two genes were most abundant in −2-cm specimens of *ppdk-1* and *dct2* leaves and in 0-cm specimens of wild type leaves (Fig. 6M and N), indicating that the suberin-related genes were turned on earlier in the mutant leaves than in wild-type leaves. When we evaluated maize genes that the MapMan tool identified as PD components, there were no significant differences in expression profiles in the mutant compared to wild-type leaves (SI Appendix, Fig. S7 and Dataset S1).

## Discussion

The timeline of chloroplast diversification in M and BS cells that we delineate included assembly of thylakoids with distinct architectures at location of ligule 2 in leaf 3, taken as the 0 cm reference point, and cell type-specific accumulation of chloroplast proteins at 4 cm. The 0- to 4-cm sector corresponded to sections in which PD across M and BS acquire sphincters and cytoplasmic sleeves. Symplastic dye movement was impeded after the structural changes of PD in a callose-dependent manner, implying that the mature PD can be gated. Furthermore, the BS cell wall became suberized to inhibit apoplastic solute movement as previously described (33). Based on these findings, we concluded that PD development at the M-BS interface overlaps with the chloroplast diversification in the two cell types.

Under conditions when the C4 cycle is disrupted, in *ppdk-1* and *dct2* mutant leaves, PD closure and suberin deposition were detected in the 0-cm zone. We hypothesize that the expedited modifications are due to a feedback reaction to aberrant concentration gradients of C4 metabolites. Given that carbon transport occurs more rapidly than carbon assimilation in C4 pathways (34), tight control of metabolite distribution is critical for coordinating enzyme reactions. In *ppdk-1* or *dct2* leaf cells, abnormal balance of C4 metabolites may stimulate a regulatory mechanism that alters PD transport. However, we did not observe a shift in the timing of cell type-specific chloroplast protein expression in the mutant leaves (SI Appendix. Fig. S5). It is possible that the genetic program for constructing dimorphic chloroplasts may be less sensitive or slower in reacting to the trouble in the mutants.

Our TEM and ET approaches yielded results that agree with previous structural, transcriptomic, and proteomic studies of the maize C4 leaf development. Using cell-type specific RNA-seq, Tausta *et al*. (2014) demonstrated that genes involved in the C4 cycle and PSII-related genes begin to be differentially expressed in M cells and BS cells from the sink to source transition zone (35). The transition zone described by Tausta *et al*. is close to our 0-cm position, where we discerned varying ultrastructural features in chloroplasts. The 0-cm position equals to the zone where light availability changes. Below the position, leaf 3 cells are wrapped inside the older leaves. Above the position, leaf 3 cells are exposed to light and are capable of photosynthesis. In a detailed proteomic analysis of maize BS cells and vascular bundle samples, normalized levels of PSII subunits decreased in samples 4.5 cm from the leaf 3 ligule, suggesting that cyclic electron transport becomes dominant in the bundle sheath at this location (13). This location overlaps with the 0-cm zone in our classification, and we found that grana stacks were scarcer in BS chloroplasts than in M chloroplasts in this section. In TEM and ET images, we detected structural dimorphism of the M and BS chloroplasts in 0-cm sections. Fluorescence microscopy-based protein localization methods failed to distinguish M and BS chloroplasts in 0-cm sections, however; probably because differences in protein compositions were too small for detection by immunolabeling.

Accelerated PD development in the mutant lines does not appear to be a consequence of seedling lethality. Their homozygous seedlings did not exhibit developmental defects except that their leaves were pale green. Leaf formation and tissue organization in homozygous genotypes were identical to those in wild-type seedlings. Using immunofluorescence microscopy analyses, we demonstrated that the biogenesis of two chloroplast types was not inhibited in the mutants. However, mutant BS chloroplasts in the 4- and 10-cm sections had less amounts of starch, suggesting that their carbon fixation is less efficient than wild type BS chloroplasts. It was previously shown that photosynthetic genes and genes encoding factors necessary for the C4 cycle are downregulated in the mutant leaves (7, 8), although gene expression patterns were largely conserved among wild type, *ppdk-1*, and *dct2* mutant leaves (SI Appendix, Fig. S4).

PD structures and permeability evolve as the cells that they connect undergo differentiation (36, 37). PD transport is regulated to coordinate organ development and tissue patterning, because non-cell autonomous signaling molecules and transcription factors diffuse through PD (38–40). A recent ET study showed that nascent PD in *Arabidopsis* root meristem cells lack cytoplasmic sleeves, but sleeves do form in PD of columella cells derived from meristem cells (41). Contrary to the idea that molecular transport happens through the cytoplasmic sleeves, CFDA and GFP diffused among meristem cells despite the finding that PD in these cells lack gaps. Our connectivity assay data are consistent with the observation. CFDA crossed the M-BS boundary in the 0-cm leaf samples, where cytoplasmic sleeves were not evident.

Sphincter modules have been reported in PD, and they are linked to inhibition of cell-to-cell transport. In dormant shoot apical cells of birch trees, symplastic transport is blocked, and PD of the inactive cells are clogged with electron-dense collars (42). When the cells are activated, the collars disappear, and amounts of callose at the PD are reduced. In the maize leaf, sphincters in PD spanning the M-BS cell wall are associated with slow photosynthesis in plants grown at low temperature (43). We demonstrated that sphincters are larger in the *ppdk-1* mutant leaves and that inhibition of callose synthase facilitated CFDA movement. Because sphincter and callose can obstruct cytoplasmic sleeves, we speculate that sleeves open or close PD at the M-BS boundary, regulated by upstream signals. Another possibility is that the sphincter may stabilize PD pores when suberin infiltrates the BS cell wall.

## Materials and Methods

*Zea mays* B73 (wild type for *ppdk-1, ppdk-2*, and *ppdk-3*), W22 (wild type for *dct2*) and five mutant lines (*ppdk-1, dct2, ppdk-2*, and *ppdk-3*) were grown from seeds in a growth chamber with 12 h:12 h light/dark and 31 °C light/22 °C dark cycle with light intensity of 550 μmol/m^2^/sec and relative humidity of 50% as previously described (9). Four segments of maize leaf tissues were collected from leaf 3 on day 10 after planting, base (0 cm, the position of leaf 2 ligule as the 0 point), early stage (−4 cm, 4 cm below the 0 point), maturing stage (4 cm, 4 cm above the 0 point) and mature stage (10 cm; 10 cm above the 0 point, close to the leaf tip). This sampling scheme was adapted from Li *et al*. (2010). Leaf tissues were pooled from 10 seedlings for each genotype and each stage. At least three specimens from each pool were examined for morphometric comparison as well as protein localization experiments. The *dct2* mutant line was isolated from the UniformMu population (https://www.maizegdb.org/uniformmu) and backcrossed 4 generations to W22. Leaf samples for microscopy analyses were harvested within 2-4 h after the onset of the light period. For TEM imaging of starch particles, however, leaf samples were dissected at 2 h before the end of the light period. Other methods are described in *SI Appendix, Materials and Methods*.

## Acknowledgments

We are grateful for support from the Hong Kong Research Grant Council (GRF14126116 GRF14121019, AoE/M-05/12, C4002-17G), the Rural Development Administration of Korea (Project No. 10953092019), and the Chinese University of Hong Kong (Direct Grants). We thank Drs. Bill Lucas (University of California, Davis), Silin Zhong (Chinese University of Hong Kong), Inhwan Hwang (Pohang University of Science and Technology, South Korea), and Sascha Offermann (Leibniz Universität Hannover) for their helpful comments in designing experiments.

## References

1. Leegood RC (2002) C4 photosynthesis: principles of CO_2_ concentration and prospects for its introduction into C3 plants. J Exp Bot 53(369):581–590.

2. Taniguchi M, Cousins AB (2018) Significance of C4 Leaf Structure at the Tissue and Cellular Levels. The Leaf: a Platform for Performing Photosynthesis, The Leaf: A Platform for Performing Photosynthesis. eds Adams WW III, Terashima I (Springer International Publishing, Cham), pp 255–279.

3. Edwards GE, et al. (2001) Compartmentation of photosynthesis in cells and tissues of C(4) plants. J Exp Bot 52(356):577–590.

4. Lundgren MR, Osborne CP, Christin P-A (2014) Deconstructing Kranz anatomy to understand C4 evolution. J Exp Bot 65(13):3357–3369.

5. Sedelnikova OV, Hughes TE, Langdale JA (2018) Understanding the Genetic Basis of C 4Kranz Anatomy with a View to Engineering C 3Crops. Annual review of genetics 52(1):249–270.

6. Rao X, Dixon RA (2016) The Differences between NAD-ME and NADP-ME Subtypes of C4 Photosynthesis: More than Decarboxylating Enzymes. Front Plant Sci 7:805–9.

7. Weissmann S, et al. (2016) Interactions of C 4Subtype Metabolic Activities and Transport in Maize Are Revealed through the Characterization of DCT2Mutants. Plant Cell 28(2):466–484.

8. Zhang Y, et al. (2018) Characterization of maize leaf pyruvate orthophosphate dikinase using high throughput sequencing. J Integr Plant Biol 60(8):670–690.

9. Li P, et al. (2010) The developmental dynamics of the maize leaf transcriptome. Nat Genet 42(12):1060–1067.

10. Pribil M, Labs M, Leister D (2014) Structure and dynamics of thylakoids in land plants. J Exp Bot 65(8):1955–1972.

11. Mai KKK, et al. (2019) Electron Tomography Analysis of Thylakoid Assembly and Fission in Chloroplasts of a Single-Cell C4 plant, Bienertia sinuspersici. Sci Rep 9(1):19640.

12. Liang Z, et al. (2018) Thylakoid-Bound Polysomes and a Dynamin-Related Protein, FZL, Mediate Critical Stages of the Linear Chloroplast Biogenesis Program in Greening Arabidopsis Cotyledons. THE PLANT CELL ONLINE 30(7):1476–1495.

13. Majeran W, et al. (2010) Structural and metabolic transitions of C4 leaf development and differentiation defined by microscopy and quantitative proteomics in maize. THE PLANT CELL ONLINE 22(11):3509–3542.

14. Faulkner C (2018) Plasmodesmata and the symplast. Current Biology 28(24):R1374–R1378.

15. Ueki S, Citovsky V (2011) To Gate, or Not to Gate: Regulatory Mechanisms for Intercellular Protein Transport and Virus Movement in Plants. Molecular Plant 4(5):782–793.

16. Deinum EE, Mulder BM, Benitez-Alfonso Y (2019) From plasmodesma geometry to effective symplasmic permeability through biophysical modelling. eLife 8. doi:10.7554/eLife.49000.

17. Brunkard JO, Runkel AM, Zambryski PC (2015) The cytosol must flow: intercellular transport through plasmodesmata. Curr Opin Cell Biol 35:13–20.

18. Han X, et al. (2014) Auxin-callose-mediated plasmodesmal gating is essential for tropic auxin gradient formation and signaling. Dev Cell 28(2): 132–146.

19. Koh E-J, et al. (2011) Callose deposition in the phloem plasmodesmata and inhibition of phloem transport in citrus leaves infected with “Candidatus Liberibacter asiaticus.” Protoplasma 249(3):687–697.

20. Zavaliev R, Ueki S, Epel BL, Citovsky V (2010) Biology of callose (β-1,3-glucan) turnover at plasmodesmata. Protoplasma 248(1): 117–130.

21. Danila FR, Quick WP, White RG, Caemmerer S, Furbank RT (2019) Response of plasmodesmata formation in leaves of C 4grasses to growth irradiance. Plant Cell and Environment 42(8):2482–2494.

22. Hatch MD (1987) C4 photosynthesis: a unique blend of modified biochemistry, anatomy and ultrastructure. Biochim Biophys Acta 895:81–106.

23. Sharpe RM, Offermann S (2013) One decade after the discovery of single-cell C4 species in terrestrial plants: what did we learn about the minimal requirements of C4 photosynthesis? Photosynthesis Research 119(1-2):169–180.

24. Ludwig M (2016) The Roles of Organic Acids in C4 Photosynthesis. Front Plant Sci 7:251–11.

25. Arrivault S, et al. (2017) Metabolite pools and carbon flow during C 4photosynthesis in maize: 13CO 2labeling kinetics and cell type fractionation. J Exp Bot 68(2):283–298.

26. Mertz RA, Brutnell TP (2014) Bundle sheath suberization in grass leaves: multiple barriers to characterization. J Exp Bot 65(13):3371–3380.

27. Danila FR, Quick WP, White RG, Furbank RT, Caemmerer von S (2016) The Metabolite Pathway between Bundle Sheath and Mesophyll: Quantification of Plasmodesmata in Leaves of C3 and C4 Monocots. THE PLANT CELL ONLINE 28(6):1461–1471.

28. Evert RF, Eschrich W, Heyser W (1977) Distribution and structure of the plasmodesmata in mesophyll and bundle-sheath cells of Zea mays L. Planta 136(1):77–89.

29. Evert RF, Russin WA, Bosabalidis AM (1996) Anatomical and ultrastructural changes associated with sink-to-source transition in developing maize leaves. International Journal of Plant Sciences 157(3):247–261.

30. Daum B, Kuhlbrandt W (2011) Electron tomography of plant thylakoid membranes. J Exp Bot 62(7):2393–2402.

31. White RG (2012) Using chlorophyll fluorescence to rapidly discriminate C3 from C4 photosynthesis in plants (Manchester, UK), European Microscopy Congress poster# LS2.8.

32. Pfündel E, Neubohn B (1999) Assessing photosystem I and II distribution in leaves from C4 plants using confocal laser scanning microscopy. Plant Cell and Environment 22(12):1569–1577.

33. Laetsch WM (1974) The C4 syndrome: a structural analysis. Annual Review of Plant Physiology 25:27–52.

34. Laisk A, Edwards GE (2000) A mathematical model of C_4_ photosynthesis: The mechanism of concentrating CO_2_ in NADP-malic enzyme type species. Photosynthesis Research 66(3):199–224.

35. Tausta SL, et al. (2014) Developmental dynamics of Kranz cell transcriptional specificity in maize leaf reveals early onset of C4-related processes. J Exp Bot 65(13):3543–3555.

36. Nicolas WJ, Grison MS, Bayer EM (2017) Shaping intercellular channels of plasmodesmata: the structure-to-function missing link. J Exp Bot 69(1):91–103.

37. Roberts AG, Oparka KJ (2003) Plasmodesmata and the control of symplastic transport. Plant Cell and Environment 26(1):103–124.

38. Kitagawa M, Jackson D (2019) Control of Meristem Size. Annual Review of Plant Biology 70(1):269–291.

39. Otero S, Helariutta Y, Benitez-Alfonso Y (2016) Symplastic communication in organ formation and tissue patterning. Current Opinion in Plant Biology 29:21–28.

40. Ham B-K, Li G, Kang B-H, Zeng F, Lucas WJ (2012) Overexpression of Arabidopsis Plasmodesmata Germin-Like Proteins Disrupts Root Growth and Development. THE PLANT CELL ONLINE 24(9):3630–3648.

41. Nicolas WJ, et al. (2017) Architecture and permeability of post-cytokinesis plasmodesmata lacking cytoplasmic sleeves. Nature Plants 3(7): 1–11.

42. Rinne PL, Kaikuranta PM, Van Der Schoot C (2001) The shoot apical meristem restores its symplasmic organization during chilling-induced release from dormancy. Plant J 26(3):249–264.

43. Bilska A, Sowinski P (2010) Closure of plasmodesmata in maize (Zea mays) at low temperature: a new mechanism for inhibition of photosynthesis. Annals of Botany 106(5):675–686.

