## Supplementary information for "Changes in the plasmodesma structure and permeability at the bundle sheath and mesophyll interface during the maize C4 leaf development"

**This PDF file includes:**

Supplementary Information Text

Figures S1 to S7

Legends for Datasets S1 and Table S1

SI References

**Supplementary Text**

**Materials and Methods**

**Transmission electron microscopy (TEM) and electron tomography**

TEM and electron tomography analyses of PD were carried out as described in Kang (1) and Koh et al. (2). We examined maize leaf samples preserved by high-pressure freezing for electron tomographic characterization of three-dimensional thylakoid structures. However, we employed chemically fixed samples for determining PD structures and starch particle analysis. We had to capture TEM micrographs from more than hundreds of PD and chloroplasts for statistical comparisons of PD and chloroplasts in multiple stages from several genetic backgrounds. It was impossible to find sufficient numbers of intact PD or chloroplasts in maize leaf samples processed by HPF because of freezing damages.

Maize leaves (leaf 3) were dissected and cryofixed with an HPM100 machine (Leica Microsystems). The frozen samples were freeze substituted in 2% OsO4 in anhydrous acetone (Leica EM AFS2) at -80 °C for 24 hrs and slowly warmed from – 80 °C to -45°C over 36 hrs. After brought to room temperature, the samples were washed with anhydrous acetone three times, and embedded in Embed-812 resin (Electron Microscopy Sciences; Cat. No. 14120) and cured at 60 °C for 24 hrs. For chemical fixation, maize leaf segments were dissected in 2.5% glutaraldehyde in 0.1 M cacodylate buffer (pH 7.4) and incubated in the fixative solution overnight at 4°C. After thorough washing (4-5 times) with 0.1 M cacodylate buffer (pH 7.4), the samples were post-fixed in 1% $OsO_4$ in deionized water for 2 hrs at room temperature. Excess $OsO_4$ was removed by washing with water (4-5 times), and the samples were dehydrated with a graduated acetone series in deionized water (from 10% to 100% acetone). Finally, they were embedded in Embed-812 resin (Electron Microscopy Sciences, Cat. No. 14120) and polymerized at 60 °C for 24 hrs. Ultrathin sections (90-nm thick) were collected on formvar-coated copper slot grids (Electron Microscopy Sciences; Cat. No. GS2010-Cu) and the sections were post-stained with uranyl acetate and lead citrate solutions. The sections were examined with a Hitachi 7650 TEM (Hitachi-High Technologies) operated at 80 kV.

Chloroplast sizes and starch particle abundances (SI Appendix, Fig. S6) were quantified from TEM micrographs. After loading micrographs in ImageJ (ver. 1.53d), outlines were drawn around chloroplasts and starch particles in chloroplasts. Areas inside the outlines were acquired with the “Measure” command under the “Analyze” tap. The values were transferred to Microsoft Excel (ver. 16.40). for calculate their ratios, average ratios, and their standard deviations. 30-40 chloroplasts in 20-30 M and BS cells (from 3 plants) were examined for each ratio calculation.

A series of semi-thick sections (250 nm) were collected on copper grids. After post-staining and gold particle coating, tilt series were collected with a 200-kV Tecnai F20 intermediate voltage electron microscope (+60° to -60° at an interval of 1.5° interval around two orthogonal axes). Tomogram calculation and 3D model rendering were performed with the IMOD software package as described previously(3, 4).

For morphometric analysis of thylakoids, stroma lamellae were defined as thylakoids consisting of single layer, and grana thylakoids were defined as thylakoids with membranes in contact with opposing membranes to constitute a stack. The largest number of disks in a granum was assigned as the number of lamellae. For PD quantitative analyses, outer diameters of sphincter rings were measured. Widths of cytoplasmic sleeves corresponded to the average width of the two gaps from the desmotubule to the plasma membrane (right and left sides) in each PD.

**Immunofluorescence microscopy localization of chloroplast proteins**

Maize leaves were dissected in 4% paraformaldehyde and fixed under vacuum for 90 min at room temperature for 3 hrs. The fixed samples were dehydrated through a graduated ethanol series (from 10% to 100% ethanol in water) and embedded in LR White resin (Electron Microscopy Sciences, Cat. No. 14383). After curing sample blocks, semi-thick sections (400 nm) were prepared with an ultramicrotome (Leica UC7, Leica Microsystems) and collected on glass slides (SuperFrost$^{TM}$, Thermo Scientific). After blocking with 2% non-fat milk dissolved in 0.2% PBST(phosphate buffered saline with 0.2% Tween 20) for 1 hr, the sections were incubated with primary antibodies (1:400 dilution) overnight at 4 °C. Information about the PsbO antibody is in Liang et al. (2018) (5). The other three antibodies, Lhca (Cat. No. AS01 005), CURT1A (Cat. No. AS08 316), and Rubisco (RbcL, Rubisco large subunit, form I; Cat. No. AS03 037), were purchased from Agrisera (https://www.agrisera.com). The cell wall was stained with an antibody against (1-3)(1-4)-beta-glucan (Biosupplies Australia, Cat. 400-3). The sections were rinsed five times with the blocking buffer, then were probed with the fluorescently conjugated secondary antibodies (1:1000 dilution; anti-mouse Alexa Fluor 488 or anti-rabbit Alexa Fluor 568; Invitrogen) for 2 hr at room temperature. Excess stain was removed by rinsing with 0.2% PBST and deionized water. Micrographs were captured with a Carl Zeiss PALM Inverted Microscope (www.zeiss.com). Immunoblot analysis of the four proteins were carried out as explained in Liang et al. (2018) (3).

For quantifying fluorescence intensities of chloroplasts, we loaded micrographs in ImageJ (ver. 1.53d), and outlines were drawn around chloroplasts to set regions of interest (ROI). Using the "Measure" command, we acquired mean intensity values inside the outlines. After collecting 30-40 mean values from randomly chosen chloroplasts in M and BS cells (from 3 plants), they were normalized to the cell wall fluorescence intensity in each micrograph. Averages and standard deviations of M or BS chloroplasts in 0, 4, and 10 cm sections were calculated with Microsoft Excel (ver. 16.40).

**Chlorophyll autofluorescence imaging**

Free-hand maize leaf cross-sections were mounted in water on glass slides (SuperFrost$^{TM}$, Thermo Scientific), and were examined using an SP8 confocal microscope to acquire the autofluorescence images of chloroplast and cell wall. After excitation with a 638 nm laser, emission wavelengths of 650-720 nm and 720-800 nm were imaged for PSII an PSI chlorophyll autofluorescence, respectively. The cell wall was visualized using its autofluorescence (excitation - 405 nm, emission - 420-480 nm).

**Starch staining**

Maize leaves (leaf 3) were dissected and cleared by incubating in 95% ethanol overnight. The samples were stained with iodine potassium iodide (IKI) solution (6) and excess IKI solution was washed off by rinsing in deionized water. Free-hand cross-sections were examined under bright-field under a Meiji Techno MT4310L biological microscope (Meiji Techno Co.) equipped with a Leica MC120 HD microscope camera.

**Carboxyfluorescein diacetate (CFDA) transport assay**

Maize leaves (leaf 3) were cut at their bases and put them upright in a beaker containing CFDA solution (50 µg/mL; Sigma-Aldrich, Cat No. 21879-25MG-F) to feed the dye from their cut ends. After 1 h in the dark, three leaf segments (0 cm, 4 cm, and 10 cm) were collected by free-hand dissection. CFDA dye distribution in the leaf cross-sections was examined with a Leica TCS SP8 confocal laser-scanning microscope. We could not assess CFDA in BS-M pairs of -4 cm cross-sections (close to the cut end) consistently because CFDA infiltrated into the apoplastic space and overstained all cell types. For DDG treatment, leaf 3 samples were first put in a DDG solution (0.1 mM; Sigma-Aldrich, Cat. No. D8375-1G) for 0.5 h and then transferred to the CFDA solution supplemented with DDG (50 µg/mL of CFDA, 0.1 mM of DDG) for another 1 h in the dark. Leaf sections were imaged with excitation of 488 nm and an emission range of 493 nm - 555 nm for CFDA fluorescence. The cell wall was visualized from its autofluorescence (405 nm excitation and detection window of 410 nm - 483 nm emission). The CDFA transport assay was repeated three times. We drew outlines of M and BS cells in ImageJ (ver. 1.53d) to obtain CFDA intensity in the cells. M/BS ratios were calculated for 20-30 cell pairs for 0-, 4-, and 10-cm samples from 3 plants for each genotype. Average ratio values, and their standard deviations were calculated with Microsoft Excel (ver. 16.40)

**Cell wall suberin staining**

Free-hand cross-sections of maize leaves (leaf 3) were incubated with freshly prepared solution of 0.1% (w/v) berberine hemisulfate (≥95%; Sigma-Aldrich, Cat. No. B3412-10G) in lactic acid (≥85%; Sigma-Aldrich, Cat. No. 252476-500G) with 1 g/mL chloral hydrate (≥99%; Sangon Biotech, Cat. No. A600288-0250) in a Petri dish covered with aluminum foil at 65 °C for 2 hrs. The solution was removed using a pipette, and sections were thoroughly washed with distilled water at least three times. After removing the water and drying on filter paper, we transferred the sections to 0.5% (w/v) aniline blue (C.I. 42755, Water-soluble; Polysciences, Cat. No. 02570-25) in distilled water for 1 hr in the dark at room temperature. The sections were subsequently washed with distilled water three time. After mounting the sections into a drop of water on microscope slides, they were examined with a Leica TCS SP8 confocal laser-scanning microscope with excitation at 405 nm and an emission window of 500 nm - 580 nm. The experiments were repeated three times. ~20 BS cell walls from three leaf samples were examined for each stage. The staining and clearing procedures were adapted from previous publications (7, 8).

Gene expression correlation test

Correlation of gene expression tendency in wild type B73, with *ppdk-1* or with *dct2* were assessed with publicly available datasets produced by Li et al. (2010) (9), Zhang et al. (2018) (10), and Weissman et al. (2016) (11). Raw data were downloaded from SRA(Accession codes: SRA012297, PRJNA280756, PRJNA340078) and mapped to maize AGPv4 genome using bowtie2. Cufflinks was used for calculating FPKM values and the Pearson correlation of the four sections between datasets were calculated using the R programming language. Violin plots were prepared from each comparison.

Sources of datasets:
B73 from Li et al. (2010),
*DCT2* (wild type control for dct2 study) and *dct2* from Weissman et al. (2016),
*PPDK* (wild type control for ppdk study) and *ppdk-1* from Zhang et al. (2018)

**RNA-seq analysis of suberin synthesis genes and PD component genes**

Differential expression analysis of maize genes encoding factors involved in suberin synthesis and constituents of PD were performed with published RNA-seq datasets for wild-type and *dct2* mutant leaf sections (accession number GSE67722) (11). The same analysis for *ppdk* mutant alleles was carried out with the published dataset SRP082943 (10). We downloaded the datasets from NCBI and used the RNA-seq analysis pipelines cited in the published reports. The FPKM values of suberin synthesis genes and PD component genes are listed in SI Appendix, Dataset S1.

**qRT-PCR verification of transcript levels**

Total RNA samples were isolated with RNeasy Plant Mini Kit (Qiagen; Cat. No. 74903), and the samples were reverse transcribed with QuantiNova reverse transcription kit (Qiagen; Cat. No. 205411). Three replicates of RNA extraction were performed at each stage for wild-type, *ppdk-1*, and *dct2* lines. qRT-PCR assays were conducted (denaturation at 95 °C, annealing/extension at 60 °C, 40 cycles) to estimate amounts of transcripts from the selected genes using a CFX96 real-time PCR detection system (Bio-Rad). Each reaction mix (10 µL) contained 5 µL SsoAdvanced$^{TM}$ Universal SYBR Green Supermix (Bio-Rad; Cat. No. 172-5272), 1 µL of cDNA template (75 ng/µL), 0.5 µL of forward primer (500 nM) and 0.5 µL of reverse primer (500 nM). The primer sequences for qRT-PCR are in SI Appendix Table S1.

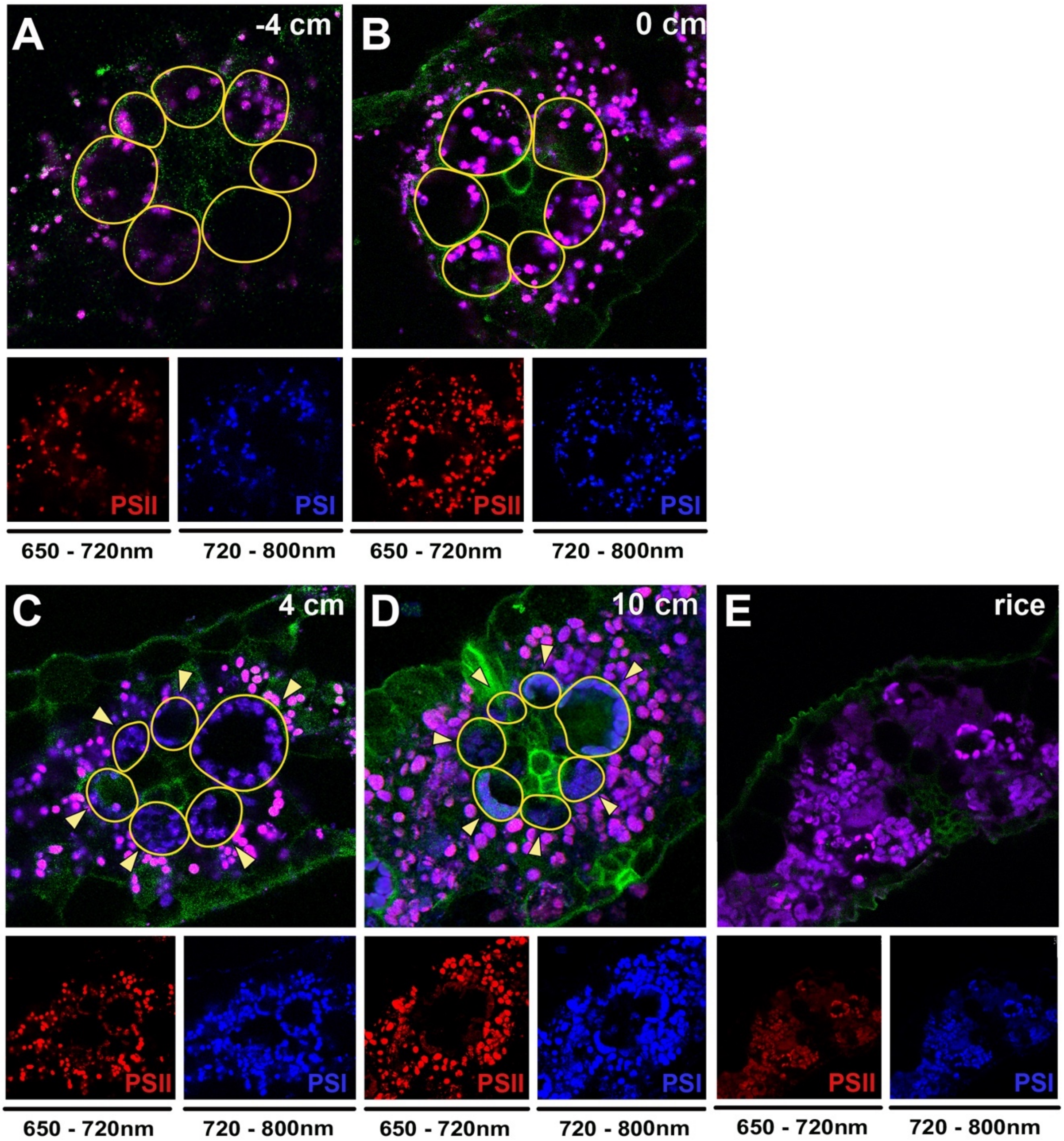


**Supplemental Fig. 1.** Distribution of photosystem (PS) II and PSI in the maize leaf.
**(A-D)** Confocal laser scanning micrographs showing cross-sections of maize leaves at -4 cm (A), 0 cm (B), 4 cm (C), and 10 cm (D). Micrographs from detection window of 650-720 nm (red, PSII) and window from 720-800 nm (blue, PSI) are provided below in each panel. Bundle sheath cells surrounding the vascular bundle are highlighted with yellow lines. In mesophyll chloroplasts with both PSI and PSII, autofluorescence from the two emission ranges overlap (magenta), whereas PSII-specific autofluorescence is reduced in bundle sheath chloroplasts in 4-cm and 10-cm sections (arrowheads in **C** and **D**). Scale bars: 25 µm.
**(E)** A rice leaf section as a control for C3 photosynthesis with a single type of chloroplasts. PSII- and PSI-fluorescence overlap (magenta).

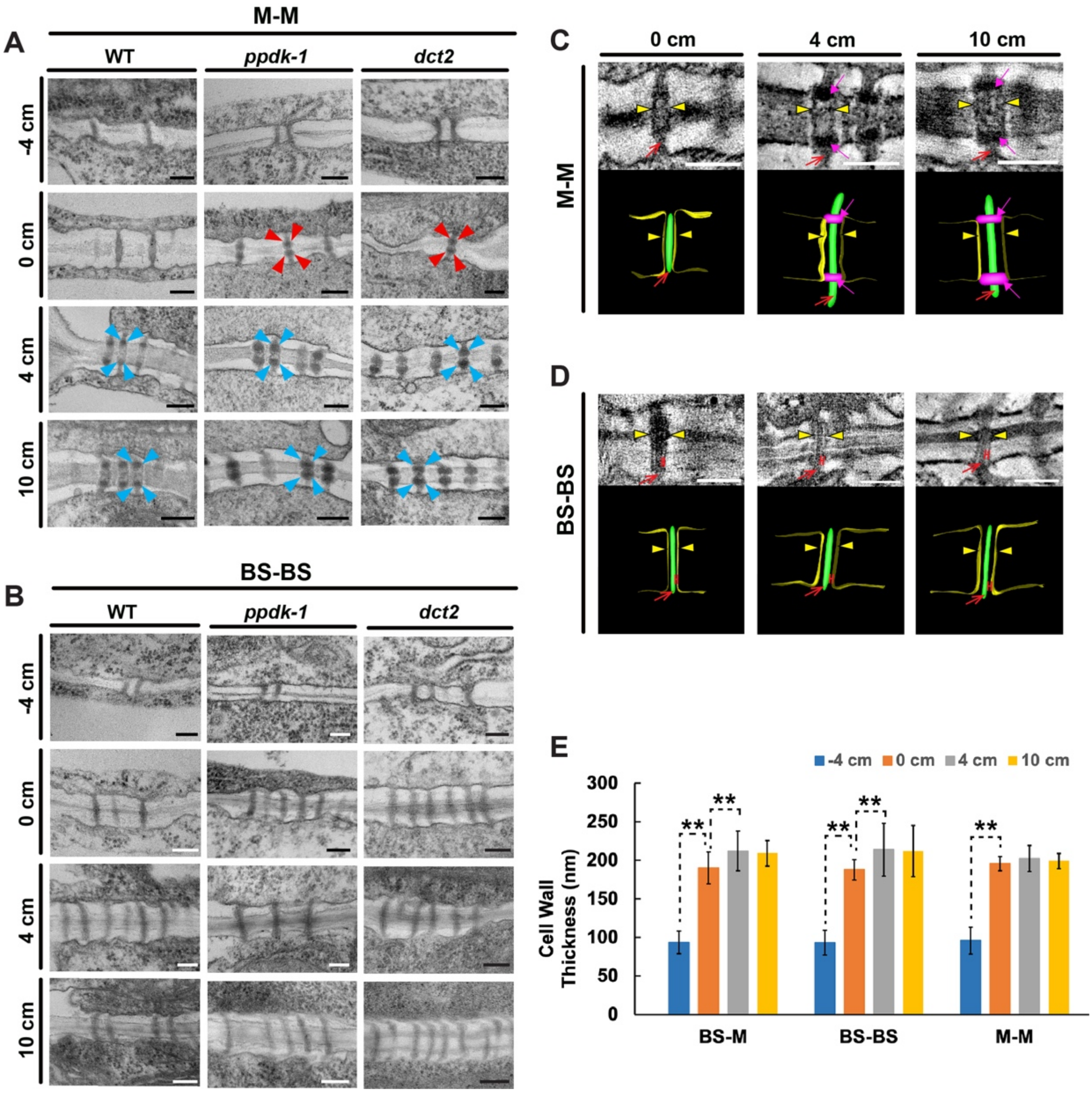


**Supplemental Fig. 2.** TEM images of PD in leaf samples of wild-type (WT) B73, *ppdk-1,* and *dct2* maize lines.

**(A and B)** TEM images of PD between mesophyll (M) cells (A) and between bundle sheath (BS) cells (B). Sphincters are marked with arrowheads. PD of 0 cm sections in the three mutant lines have sphincters (red arrowheads). Scale bars: 100 nm.

**(C and D)** Electron tomography slice images of PD and their 3D models in M-M (C) and in BS-BS (D) cell walls in WT leaves. Sphincters (magenta), desmotubules (green), plasma membrane (yellow), and cytoplasmic sleeves were marked with magenta arrows, red arrows, yellow arrowheads, and red "H"s, respectively. PD in the BS-BS cell walls lack sphincters (D). Scale bars, 150 nm.

**(E)** Thicknesses of the cell walls between BS-M, BS-BS, and M-M cells of WT leaves at the four developmental stages. Thickness were measured from TEM micrographs (-4 cm, blue bars; 0 cm, orange bars; 4 cm, gray bars; and 10 cm, dark yellow bars). Cell walls of 10 cells from three leaf samples (n=30) were measured for each bar in the graph. Error bars are standard errors (**, $p<0.01$ by Student's t-test).

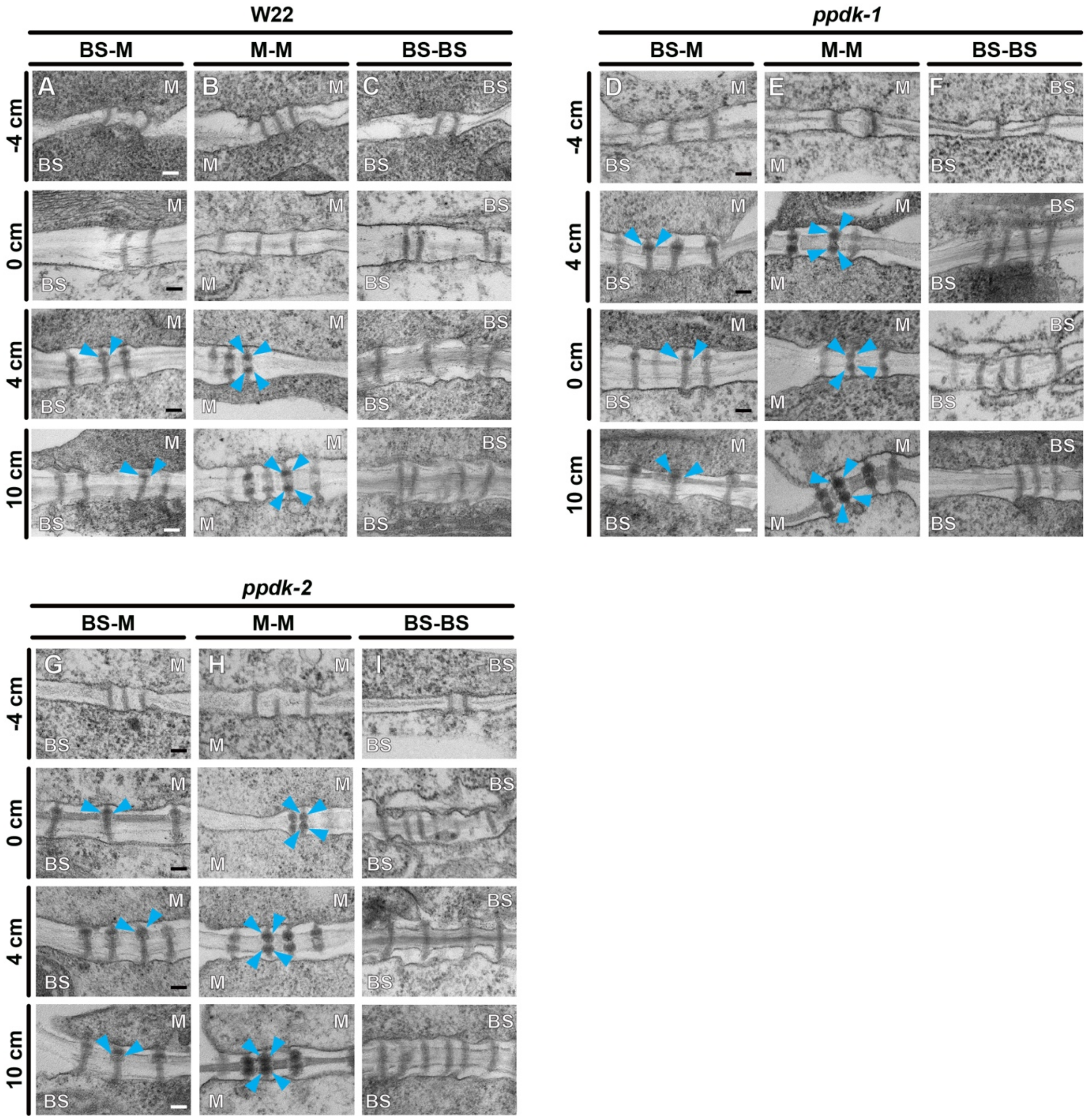


**Supplemental Fig. 3.** TEM micrographs of PD in the maize leaf samples from wild type W22 **(A-C)**, *ppdk-2* **(D-F)** *and ppdk-3***(G-I)** mutant alleles. PD in cell walls between bundle sheath (BS)-mesophyll (M), M-M, and BS-BS are shown. Sphincters are marked with blue arrowheads on the M side of the cell wall. Scale bars: 100 nm.

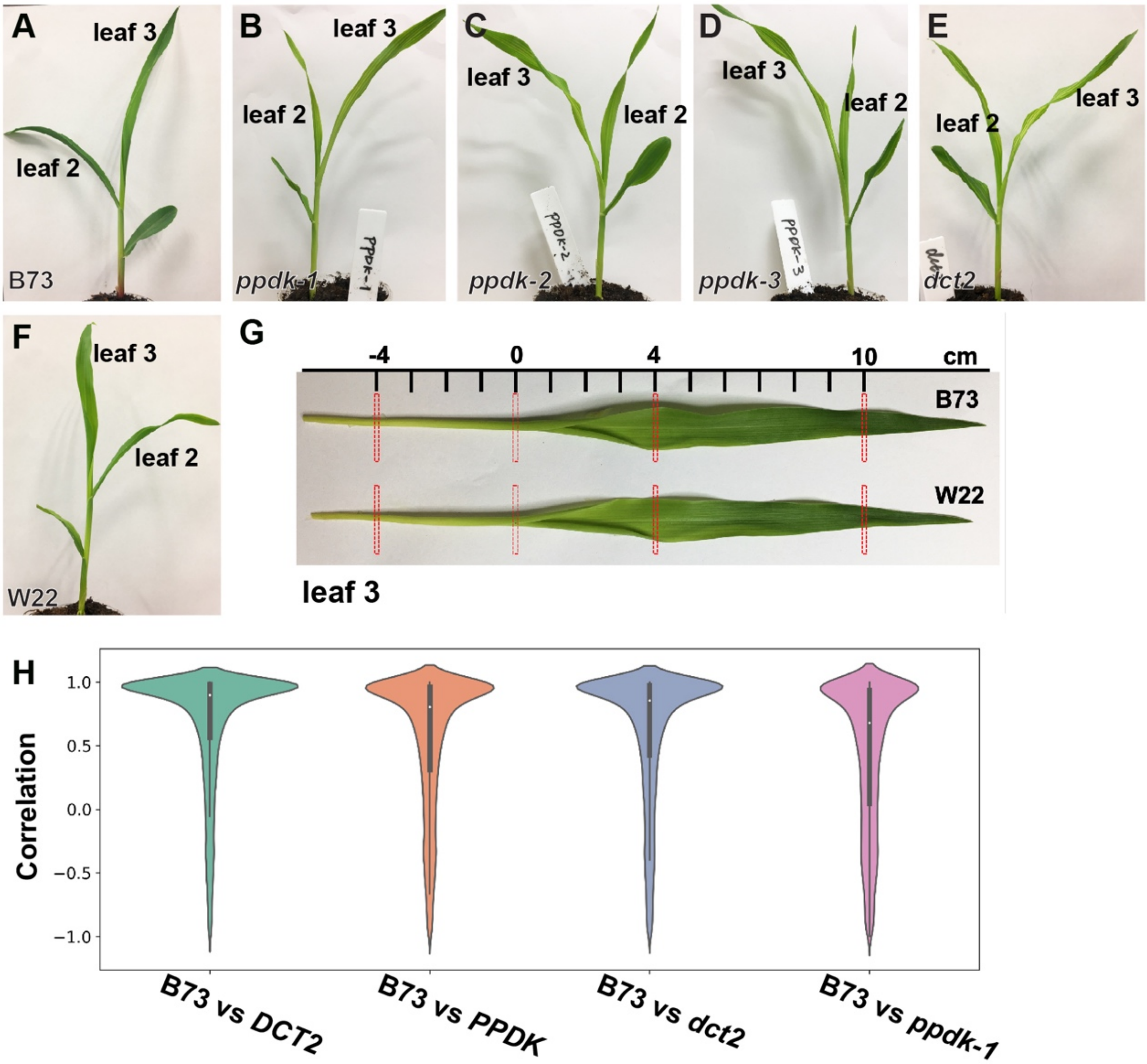


**Supplemental Fig. 4.** Photographs of 9-day-old maize seedlings and gene expression correlation in wild type and mutant leaves. **(A)** wild-type (WT) B73, **(B)** *ppdk-1*, **(C)** *ppdk-2*, **(D)** *ppdk-3*, **(E)** *dct2,* and **(F)** W22 seedlings grown under conditions described in **Methods**. Leaf 2 and leaf 3 are marked in the photos. **(G)** The four positions (-4, 0, 4, 10 cm) for developmental stages in leaf #3 from B73 and W22 inbred lines. The red rectangles mark leaf tissue samples isolated for microscopy analysis. #3 leaves from W22 are slightly shorter than those form B73. We set the 0-cm position in W22 in the same way as in B73 and isolated samples in -4, 4, and 10 cm from the 0-cm position. **(H)** The distribution of gene expression correlations in B73, wild type control for *dct2* (*DCT2*), wild type control for *ppdk-1* (*PPDK*), *dct2*, and *ppdk-1* lines. Correlation coefficients for matching genes in the datasets were illustrated in the violin plots. In general, gene expression patterns over the four leaf developmental stages in B73 are conserved among the genotypes.

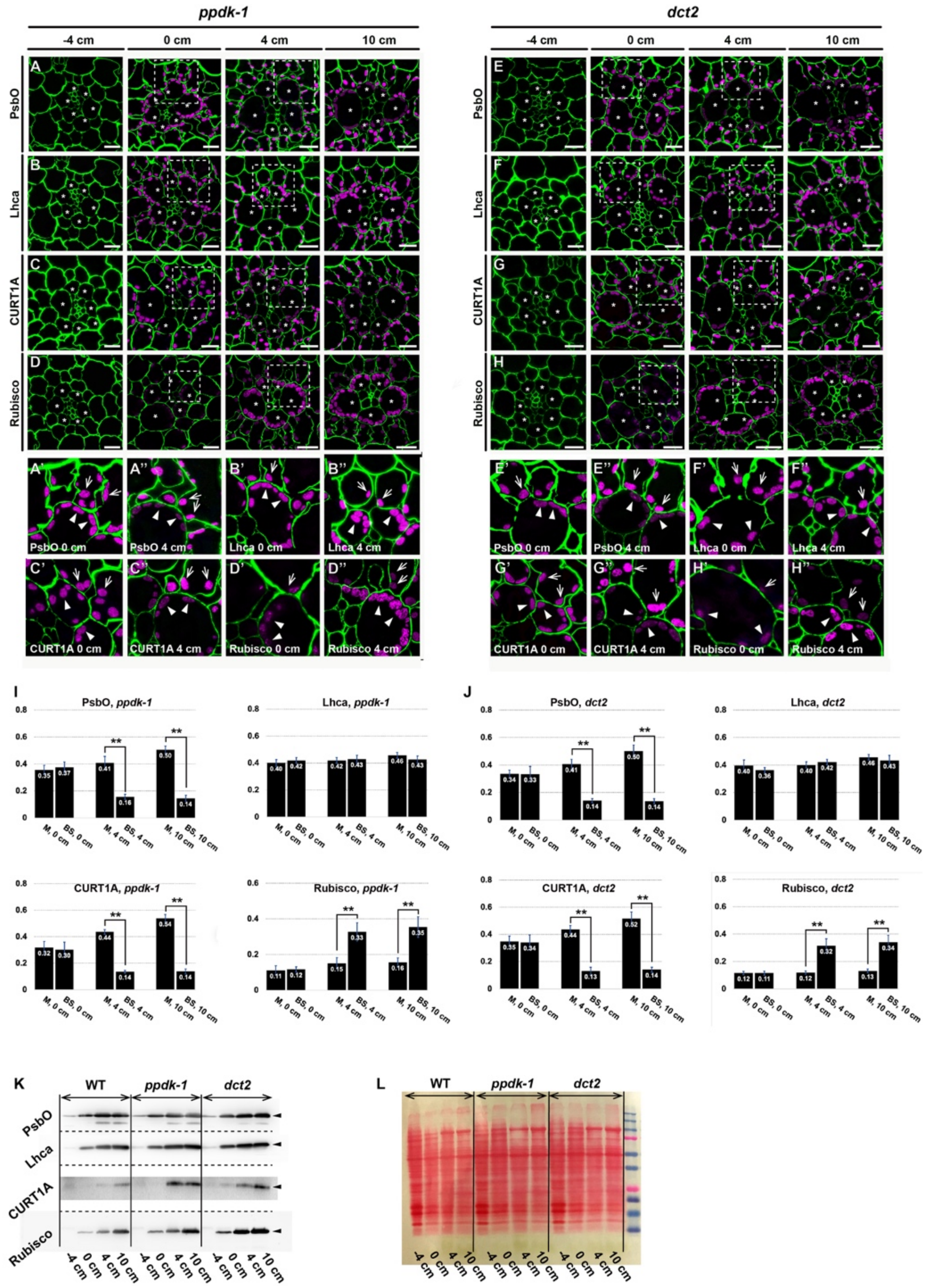


**Supplemental Fig. 5.** Immunolocalization of chloroplast proteins in mesophyll (M) and bundle sheath (BS) cells of *ppdk-1* (A-D) and *dct2* (E-H) leaves.

**(A-H)** LR white sections from four leaf locations stained for PsbO (A and E), Lhca (B and F), CURT1A (C-G), and Rubisco large subunit (D-H). The cell wall was counter-stained to illustrate Kranz anatomy (pseudo-colored in green). Higher magnification images of boxed areas in 0 and 4 cm micrographs are presented below (A'-H' and A''-H''). M and BS chloroplasts are marked with arrows and arrowheads in the magnified panels. Differential enrichment of PsbO and CURT1A in M chloroplasts was observed in 4-cm and 10-cm sections. Rubisco concentrated to BS chloroplasts in 4-cm and 10-cm sections. Lhca, a PSI subunit, accumulated evenly in the two types of chloroplasts. Scale bars: 10 µm.

**(I and J)** Average intensities were calculated from 20 randomly chosen M or BS chloroplasts in *ppdk-1* (I) and *dct2* (J) micrographs from three leaf samples. Error bars depict standard deviation (SD) (**, p<0.01 by Student t-test).

**(K and L)** Immunoblot analysis of the four proteins localized by immunofluorescence microscopy (K) and a nitrocellulose membrane stained with Ponceau S dye (L)

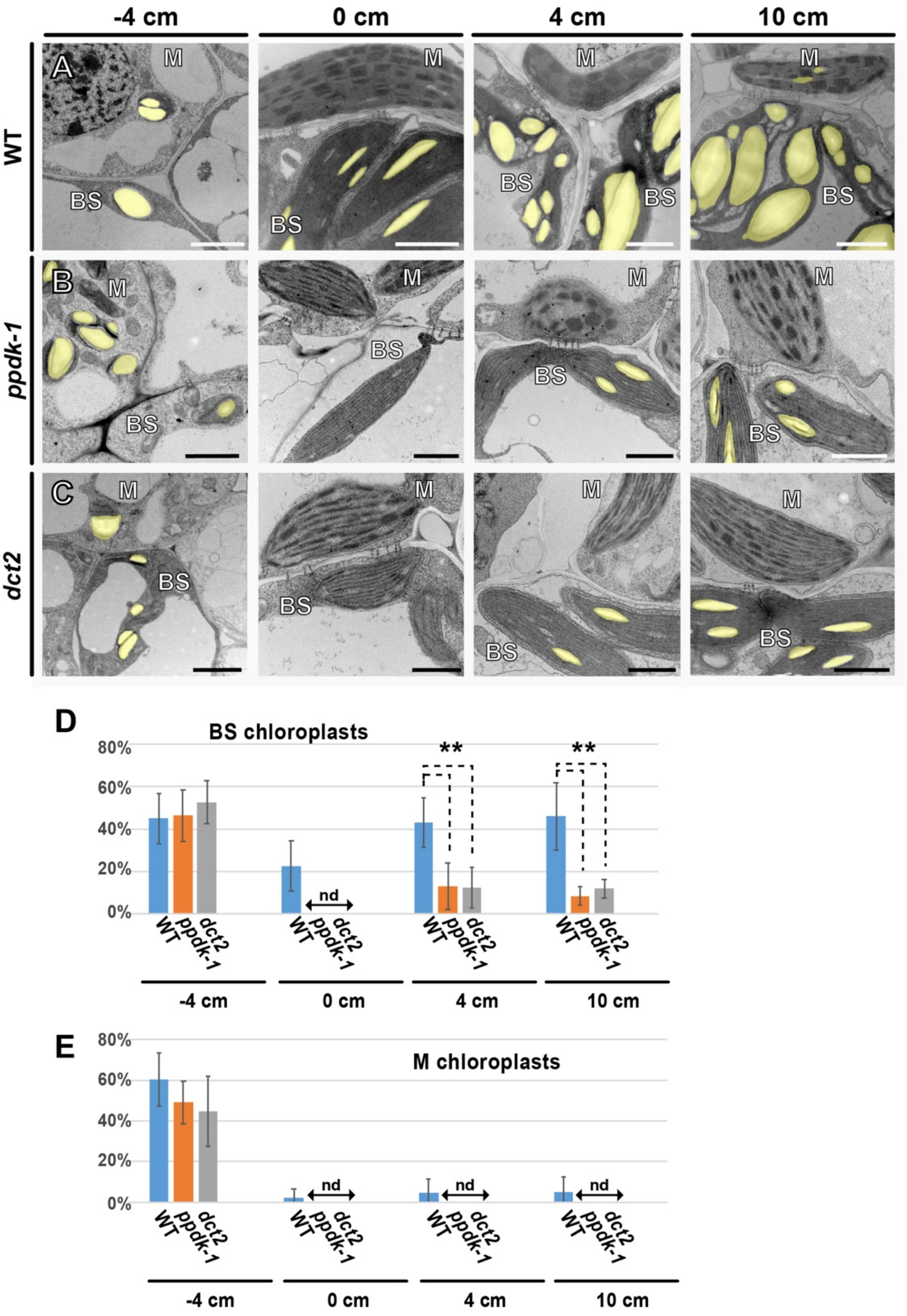


**Supplemental Fig. 6**. Starch particle accumulation in mesophyll (M) and bundle sheath (BS) chloroplasts.

**(A-D)** TEM micrographs of BS and M cells showing their chloroplasts (pseudo-colored in yellow) in wild type (WT) B73, *ppdk-1*, and *dct2* leaves. Four positions in the leaf were examined. Starch particles were pseudo-colored in yellow.

**(E-F)** Percentages of the stroma occupied by starch particles in BS **(E)** and M **(F)** chloroplasts. Chloroplast sizes and areas of starch particles were measured from TEM micrographs using ImageJ (ver. 1.53d). nd: not detected.

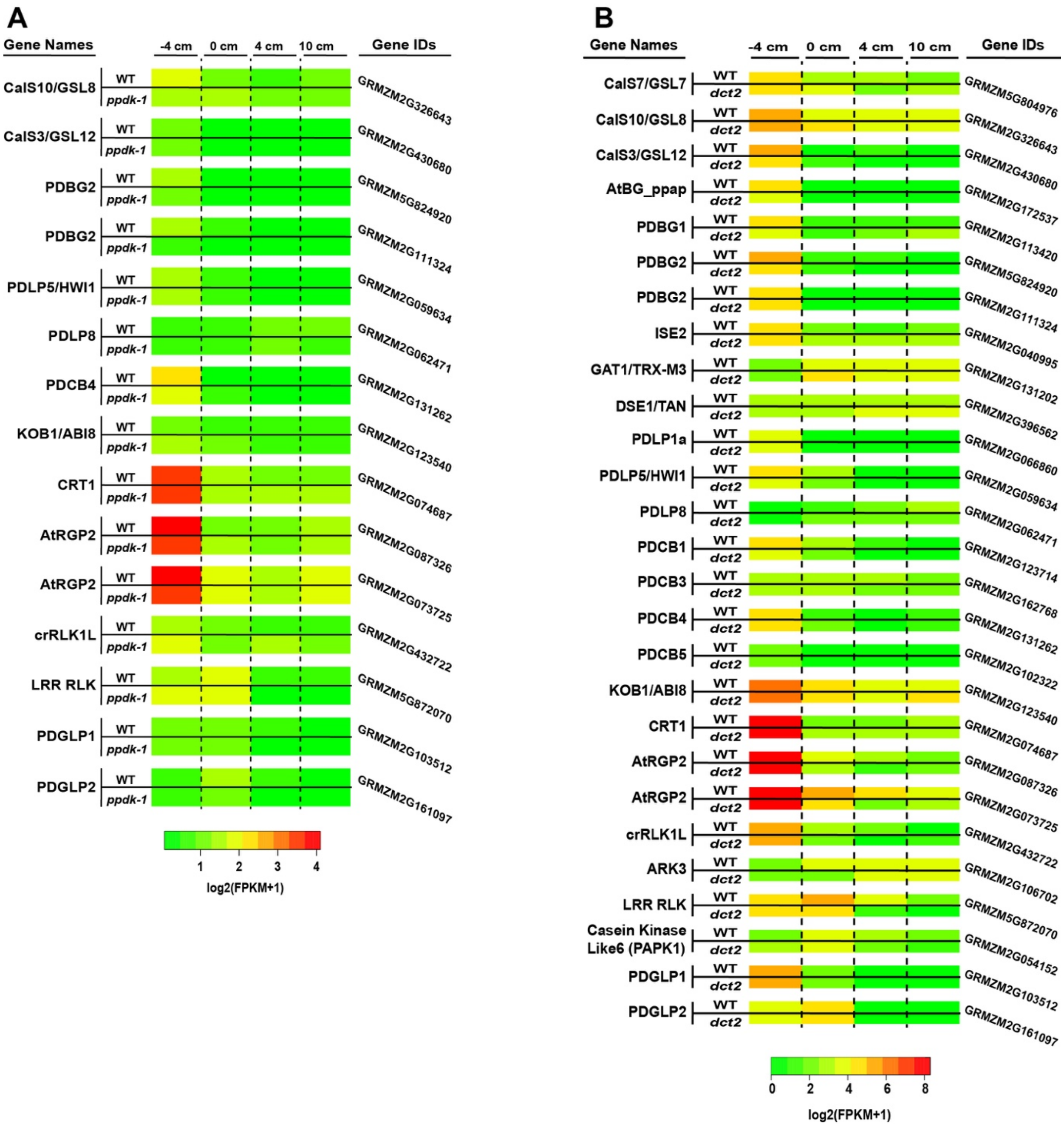


**Supplemental Fig. 7.** RNA-seq analysis of genes for PD components
**(A and B)** Heat maps showing transcript levels of genes encoding PD components in *ppdk-1* (A) and *dct2* (B) in comparison with wild type (WT). PD genes with FPKM values higher than 1 are shown (SI Appendix, Dataset. S1). None of the PD genes displayed stronger transcriptional activities in *ppdk-1* or *dct2* than in WT.

**Supplemental Dataset S1.** List of genes involved in suberin synthesis and genes encoding PD components

**Supplemental Table S1.** Primer sequences for qRT-PCR in Fig. 6

### SI References


1. Kang B (2010) Electron microscopy and high-pressure freezing of Arabidopsis. *Methods in Cell Biology* 96:259–283.

2. Koh E-J, et al. (2011) Callose deposition in the phloem plasmodesmata and inhibition of phloem transport in citrus leaves infected with "Candidatus Liberibacter asiaticus." *Protoplasma* 249(3):687–697.

3. Mai KKK, Kang B-H (2017) Semiautomatic Segmentation of Plant Golgi Stacks in Electron Tomograms Using 3dmod. *Methods Mol Biol* 1662:97–104.

4. Toyooka K, Kang B-H (2014) Reconstructing Plant Cells in 3D by Serial Section Electron Tomography. *Plant Cell Morphogenesis*, Methods in Molecular Biology. eds Zarsky V, Cvrckova F (Humana Press, Totowa, NJ, Totowa, NJ), pp 159–170.

5. Liang Z, et al. (2018) Thylakoid-Bound Polysomes and a Dynamin-Related Protein, FZL, Mediate Critical Stages of the Linear Chloroplast Biogenesis Program in Greening Arabidopsis Cotyledons. *THE PLANT CELL ONLINE* 30(7):1476–1495.

6. Ruzin SE (1999) *Plant microtechnique and microscopy* (Oxford University Press, New York).

7. LUX A, Morita S, ABE J, ITO K (2005) An Improved Method for Clearing and Staining Free-hand Sections and Whole-mount Samples*. *Annals of Botany* 96(6):989–996.

8. Yadav V, et al. (2014) ABCG Transporters Are Required for Suberin and Pollen Wall Extracellular Barriers in Arabidopsis. *THE PLANT CELL ONLINE*. doi:10.1105/tpc.114.129049.

9. Li P, et al. (2010) The developmental dynamics of the maize leaf transcriptome. *Nat Genet* 42(12):1060–1067.

10. Zhang Y, et al. (2018) Characterization of maize leaf pyruvate orthophosphate dikinase using high throughput sequencing. *J Integr Plant Biol* 60(8):670–690.

11. Weissmann S, et al. (2016) Interactions of C 4Subtype Metabolic Activities and Transport in Maize Are Revealed through the Characterization of DCT2Mutants. *Plant Cell* 28(2):466–484.
